# Hummingbird blood traits track oxygen availability across space and time

**DOI:** 10.1101/2022.01.18.476833

**Authors:** Jessie L. Williamson, Ethan B. Linck, Emil Bautista, Ashley Smiley, Jimmy A. McGuire, Robert Dudley, Christopher C. Witt

## Abstract

Predictable trait variation across environmental gradients suggests that adaptive evolution repeatedly finds the same solution to a challenge. Trait-environment associations can reflect long-term, genetic evolution across phylogenies, or short-term, plastic responses of individuals. When phylogenetic and population-level patterns match, it implies consistency between the long timescale of adaptation and the short timescale of acclimatization. Alternatively, genetic adaptation can find solutions that ‘break the rules’ of trait-environment covariation. For example, blood-hemoglobin concentration ([Hb]) increases at high elevations in animals, but genetic adaptations in some populations have been shown to augment tissue oxygenation while curtailing hemoglobin production, altering this predictable [Hb]-elevation association. Here, we tested whether species adaptation to elevation generally alters trait-environment relationships for blood. To do this, we measured blood traits of 1,217 individuals representing 77 species of Andean hummingbirds, across a 4,600 m elevational gradient. We used hierarchical Bayesian modeling to estimate blood trait responses to elevation, environmental temperature, precipitation, individual and species characteristics, and phylogeny. Strikingly, the effects of elevation on [Hb] and hematocrit (Hct) were nearly identical for individuals and species, implying that rules of elevational blood variation are set by physics of gas exchange in the hummingbird respiratory system and are unchanged by species adaptation. However, when we looked at mechanisms of [Hb] adjustment—by changes in red blood cell size or number—we did find a signal of species adaptation: To adjust [Hb], species at low and high elevations, respectively, tended to adjust cell size, whereas species at mid-elevations tended to adjust cell number. Despite scale-independent elevational variation in [Hb] and Hct, the species-specific balance of red blood cell size versus number appears to have been affected by adaptations that distinguish hummingbird species living at moderate versus high elevations.

## Introduction

Predictable variation of traits across environmental gradients can reveal both the power and limitations of adaptive evolution. Trait-environment relationships among species tend to reflect long-term, genetically fixed trait differences across phylogenies; in contrast, trait-environment relationships within species tend to reflect short-term, plastic responses of individuals with shared genetic backgrounds. When within-species and among-species patterns for a specific trait match, it implies a single, shared solution to an environmental challenge. For example, cooler waters at high latitudes are consistently associated with higher numbers of vertebrae in fish, whether comparing species or conspecific individuals (Jordan’s Rule) [1,2]. When within-species and among-species patterns conflict, it implies that genetic adaptation has found a qualitatively different solution to the same environmental challenge, altering the trait-environment relationship. The tendency for larger body sizes at colder ambient temperatures among species (Bergmann’s Rule) and within species (James’s Rule; [3]) is frequently subject to conflicting patterns at these two scales of comparison; it is detectable in only a subset of animal clades, and is reversed in others [4]. These conflicting trait patterns have been attributed to multifarious selection pressures impacting species differences in body size and thermoregulatory physiology [3], and they reflect the expected erosion of trait-environment (or trait-trait) associations over macroevolutionary time [5]. Trait-trait relationships are similarly scale-dependent [6]; for example, individual common milkweed (*Asclepia syriaca*) plants that produce higher concentrations of cardenolide toxins have reduced growth rates [7], while other species of milkweed that produce up to seven times the concentrations of cardenolide toxins as *A. syriaca* show no reduction in growth rate [8]. In this way, genetic adaptations can ‘break the rules’ that govern trait relationships. Explaining such ‘trait gradients’ across biological scales will be essential not only for understanding the functional basis of adaptation, but for predicting the functional consequence of global change for species and communities [9].

In animals, blood traits associated with oxygen transport change predictably with declining oxygen (O_2_) availability at elevation [10–13]. Most O_2_ in the blood is carried by hemoglobin (Hb) [14], the concentration of which ([Hb]) is a key determinant of blood-O_2_ carrying capacity. When lowland-adapted birds ascend to altitude, they experience reduced arterial O_2_ saturation; to compensate, they undergo erythropoiesis, which can increase [Hb] and therefore blood-O_2_ carrying capacity, enhancing O_2_ delivery to the tissues and compensating for reduced arterial O_2_ saturation [13]. In the absence of compensatory increases in plasma volume [15], increased Hb mass leads to elevated hematocrit (Hct) [16]. A suite of interrelated physical characteristics of blood—total red blood cell count (TRBC), mean cell volume (MCV), mean cell hemoglobin (MCH), and mean cell hemoglobin concentration (MCHC; Table S1)—can be modulated to optimize blood-O_2_ transport, or to compensate for other cardiac, vascular, or hematological responses to elevation. For example, MCV typically increases in acclimatized, lowland-adapted animals undergoing accelerated erythropoiesis [17].

However, genetic adaptation to high elevation can break the ‘rules’ of trait-environment covariation in blood traits. For example, evidence from high-altitude-adapted human populations and theoretical work suggests that optimal [Hb] and Hct levels may be similar to those of sea-level populations [13,18–20]. Adaptive genetic changes to hemoglobin proteins have been demonstrated in many high-altitude-native birds, including hummingbirds, House Wrens, Andean waterfowl, Bar-headed Geese, and a few dozen other bird taxa [21–25]. In hummingbirds, there is a highly predictable association between species-typical Hb β^A^13–β^A^83 genotype and elevation, with derived increases in Hb-O_2_ affinity in highland lineages and reductions in Hb-O_2_ affinity in lowland lineages [22]. Hummingbirds occurring at high altitudes exhibit two amino-acid substitutions on the β3^A^-globin subunit of Hb (Hb β3^A^-13_GLY_⟶_SER_ and Hb β3^A^-83 _GLY_⟶_SER_) that are known to affect O_2_ binding [22]. In particular, β3^A^-13_SER_ is known from only four hummingbird genera (*Patagona, Chalcostigma, Pterophanes*, and *Oreotrochilus*), all of which occur at extreme high altitudes [21,22,26]. O_2_-binding affinity of hemoglobin has critical effects on tissue oxygenation, and, in birds, binding affinity increases predictably with elevations [21,27]. It is not yet clear whether the genetic adaptations that change Hb-O_2_ binding affinity predictably affect other blood traits or their covariation with elevation. It is plausible that species-typical Hb β3^A^ genotypes associated with different O_2_-binding affinities induce predictable differences in blood traits and their reaction norms to elevation; such an association could reflect a direct functional tradeoff or an indirect effect of accrued, multi-locus adaptation to elevation at multiple loci.

The macro-physiological ‘rules’ governing blood trait variation over phylogenetic and population timescales have remained elusive because adequate comparative data are scarce. Blood traits are known to vary among avian clades (e.g., [28,29]) and exhibit elevational variation both within species (e.g., [30–32]) and among species (e.g., [30,31]). At least two additional recent studies have conducted broad-scale interspecific comparative analyses of avian [Hb] and Hct data, but both datasets were compiled from heterogeneous published data and lacked elevational sampling sufficient to estimate elevational effects [33,34]. Furthermore, no previous comparative analysis of hematological variation across elevation has included the six key parameters that affect blood-O_2_ carrying capacity per unit volume (Table S1). Thus, we continue to lack a general understanding of within- and among-species blood trait-environment relationships.

Examining the underlying mechanisms of adjustment for blood-oxygen carrying capacity can offer insight on trait-environment relationships. [Hb] is equivalent to O_2_-carrying capacity per unit volume of blood, and it can be adjusted via adjustments to cell number (TRBC), cell size (MCV), within-cell [Hb] (MCHC), or plasma volume. Cell number and cell size are both measurable components that vary among individuals and species, are subject to plastic and evolutionary adjustments, and have strong direct effects on [Hb]. Cell number and cell size are reciprocally constrained by physical space available in the circulatory system, such that individual birds can have a few large erythrocytes, many small erythrocytes, or an intermediate state for both traits [35,36]. MCHC tends to be relatively invariant across the vertebrate tree of life [35], with Hb typically comprising ∼1/3 of erythrocytes. Plasma volume adjustments can decouple Hb mass from [Hb] and, although plasma volume appears variable among human populations [15], we currently have insufficient data on elevational variation in avian plasma volume to assess its importance. Quantifying the relative contributions of TRBC and MCV in driving [Hb] variation within and among species may show that species use different mechanisms of adjusting blood-O_2_ carrying capacity, revealing signatures of adaptation that distinguish hummingbirds living at moderate versus high elevations.

In this study, we sought to test whether phylogenetic and population patterns match for blood-trait variation with environment. Specifically, we asked: Does species adaptation change trait-environment relationships? To address this question, we spent 2006–2020 collecting specimen-vouchered hematological data from 1,217 wild hummingbirds of 77 species, representing all 9 clades, and spanning ∼4,600 meters in elevation. Using these data, we then constructed hierarchical Bayesian models to estimate the responses of six blood traits to elevation, climate, individual and species characteristics, and phylogeny. The hummingbirds are a potential model clade in which to study rules of hematological variation due to their documented evolutionary genetic responses to shifts in *P*O_2_, high species richness in mountains, phylogenetically conserved elevational ranges, and relative uniformity of body size, clutch size, diet, mating system, and flight mode [21,22,37–39]. Notably, matching O_2_ supply and demand is an acute challenge for all hummingbirds because they exhibit the highest mass-specific metabolic rates among endotherms, and because they employ sustained hovering, one of the most energetically demanding forms of animal locomotion [38,40]. This study posits new rules of elevational blood trait variation, at least for hummingbirds, providing new insights into the origins and maintenance of trait-environment relationships.

## Results and Discussion

We quantified three primary ([Hb], Hct, and TRBC) and three secondary hematological indices (MCV, MCH, and MCHC; Table S1), and separately analyzed two datasets: an individual-level dataset, wherein each row contained data from one individual hummingbird (*n*=1,217), and a species-level dataset, wherein each row contained mean blood values calculated for each species, with a minimum of 2 individuals sampled (*n*=58–65 species, depending on blood trait modeled). In the individual-level data, mean [Hb] was 18.84 g/dl (±1.65), mean Hct was 58.54% (±5.15), mean TRBC was 6.37 RBC × 10^6^/mm^3^ or 6,370,000 cells per µl (±1.16), mean MCV was 93.36 fl (±17.1), mean MCH was 30.4 pg (±5.64), and mean MCHC was 32.44 g/dl (± 2.12). Complete reference values—the first such values ever published for >90% of our study species––are reported in detail in Table S2.

### Within-species blood trait-environment variation

We used the individual-level dataset to model within species (i.e., interspecific) variation using multilevel Bayesian models. For each of six blood parameters, we estimated the effects of elevation, body mass, precipitation, air temperature, species-typical Hb β^A^13–β^A^83 genotype, as well as within-species variation in elevational position (an individual’s sampled elevation relative to its species elevational range), body mass, air temperature, and precipitation. We accounted for between-species differences by incorporating a species grouping variable. Separately, we excluded sex and wing-loading as potentially important predictors (see Methods). In total, we compared six models per set using LOOIC and WAIC (Tables S3-S4; Methods). In addition to fixed effects, models included species as a random effect, and a phylogenetically correlated random intercept (i.e., covariance matrix of phylogenetic distances among taxa). Thus, our full model in each set reflects the hypothesis that all predictors, phylogeny, and species identity influence blood characteristics.

Based on the top-performing model in each set, we found that for every 1,000-m increase in elevation, [Hb] is expected to increase by 0.39 g/dl, Hct is expected to increase by 1.42 percentage points, TRBC is expected to increase by 310,000 cells per µl, and MCH is expected to decrease by 0.48 pg. Elevation strongly predicted [Hb], Hct, TRBC, and MCH (Figs 2 A–F and 3A–F). Elevational position was also a strong predictor of Hct, indicating that individuals located higher in their species’ elevational range tended to have higher Hct (Fig 3B). [Hb], TRBC, and MCH also increased with increasing elevational position, although these relationships were subtle (Fig 3A, C, E).

**Fig 1.**
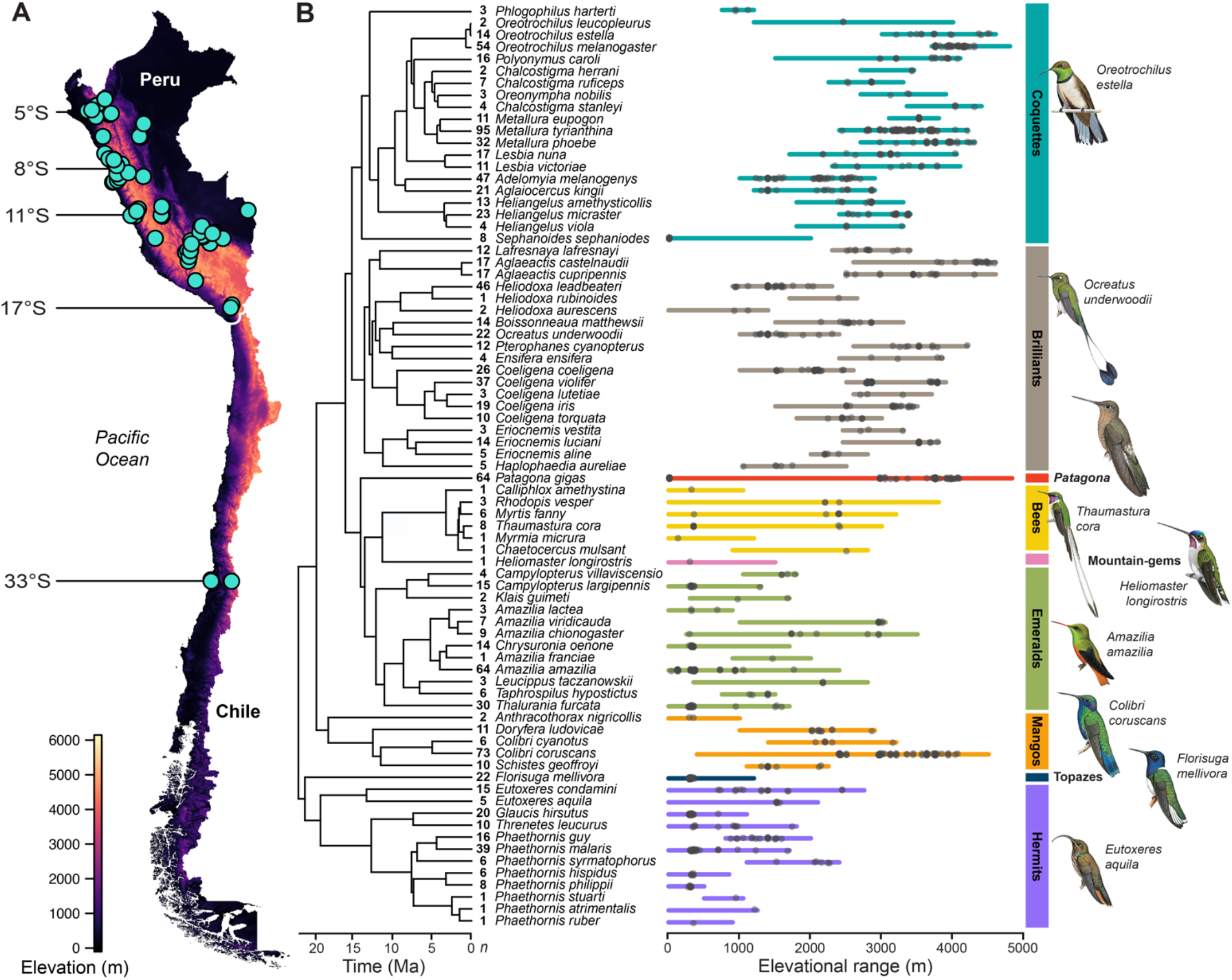
Elevational, latitudinal, and phylogenetic sampling from 2006–2020. **(A)** Sampling localities spanned 28° latitude and 4,578 meters in elevation in Peru and Chile. **(B)** We studied 77 species (‘n’ column indicates sampling depth) from all nine major hummingbird clades. Sampling (gray opaque points) is shown across each species’ elevational range (colored horizontal bars). Time-calibrated phylogeny is modified from McGuire et al. [37]. Illustrations depicting a representative from each clade are from *Birds of the World* and are reproduced courtesy of the Cornell Lab of Ornithology (©).

**Fig 2.**
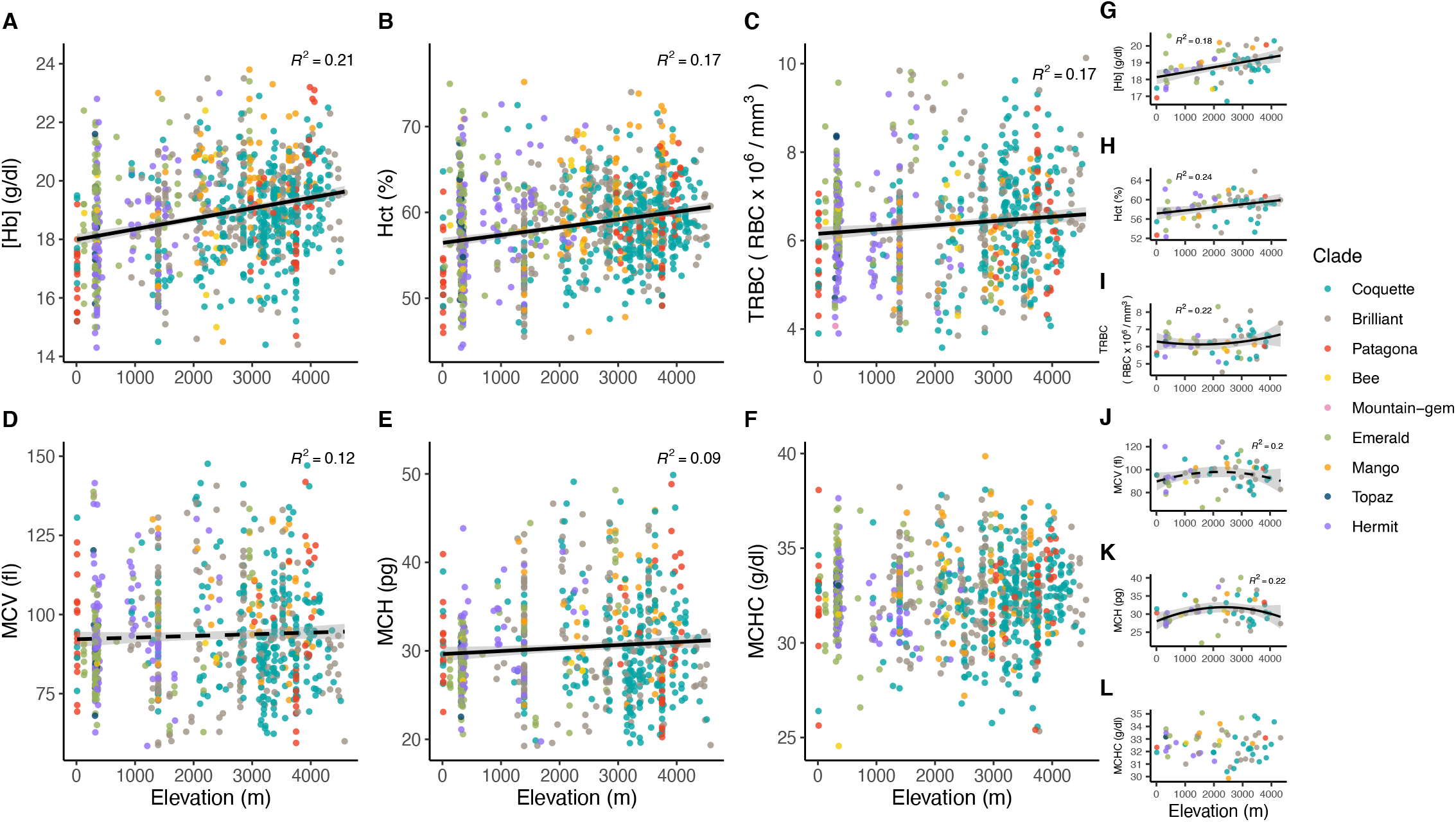
Relationship between elevation and the six studied blood parameters (hemoglobin concentration ([Hb]), hematocrit (Hct), total red blood cell count (TRBC), mean cell volume (MCV), mean cell hemoglobin content (MCH), and mean cell hemoglobin concentration (MCHC). **(A–F)** Within species comparisons, wherein each point represents a value from a single individual. **(G–L)** Among species comparisons, wherein each point represents a mean value from a single species. In all panels, solid trend lines correspond to relationships where 95% posterior credible intervals in models did not overlap zero, while dashed trend lines correspond to relationships where 89% credible intervals in models did not overlap zero (see Fig 3). Credible intervals for both elevation (at the 95% level) and *elevation*^*2*^ (at the 89% level) did not overlap zero in the TRBC model (Panel I). Colors correspond to clades in Fig 1. R^2^ values are reported from Bayesian models reported in Fig 3.

**Fig 3.**
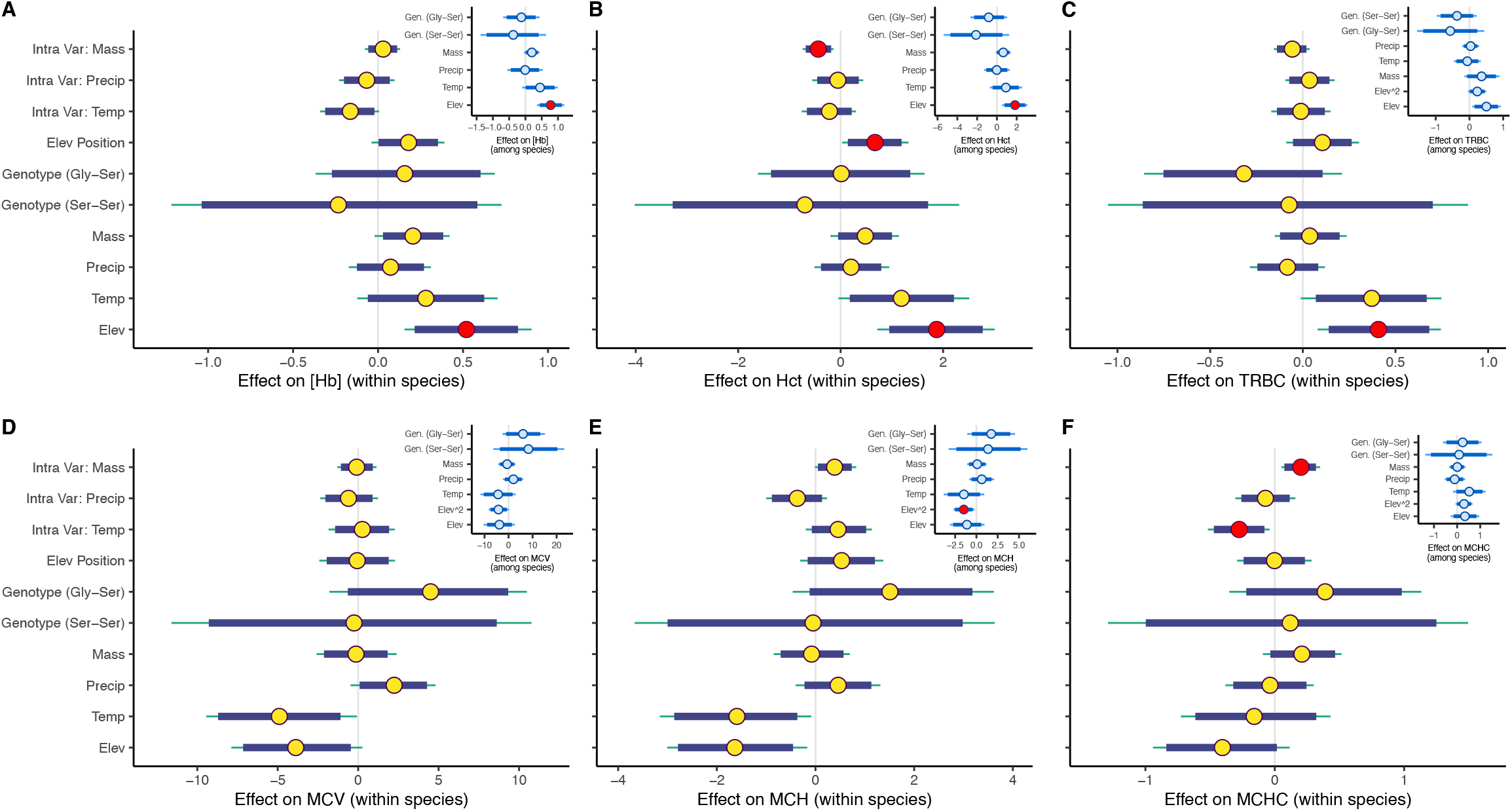
Posterior probabilities and credible intervals from full within-species (main plots) and among-species (insets) models for each of six blood traits: **(A)** hemoglobin concentration ([Hb]), **(B)** hematocrit (Hct), **(C)** total red blood cell (TRBC), **(D)** mean cell volume (MCV), **(E)** mean cell hemoglobin (MCH), **(F)** mean cell hemoglobin concentration (MCHC). Circles indicate beta estimates, thick and thin bars illustrate 89% and 95% credible intervals, respectively. Red circles denote parameters that did not overlap zero in top-ranked models. For Hb, Hct, and TRBC within-species analyses, the full model was the top model. See Tables S3-S4 for model comparison and parameter estimates from top models in other model sets.

The finding that [Hb], Hct, and TRBC increase with elevation was consistent with theoretical expectations for O_2_-carrying capacity of blood and with prior empirical findings in birds [26,30,31,33,34], humans [41,42], mammals [11,43], and other taxa [44,45]. In prior work, substantial variation in slopes of these parameters across elevation may have been attributable to variation in temporal, taxonomic, and spatial scales of comparison, as well as constraints on the time-course of evolutionary adaptation [22,42,46]. We expect increases in Hct at higher elevations because an increase in the total mass of Hb enhances oxygenation under lower ambient *P*O_2_ [18]; unless there is a proportional increase in plasma volume or MCHC, an increase in Hb mass is expected to be accompanied by an increase in the proportion of blood comprised of red blood cells. TRBC also increased with elevation and was, along with MCV, one of two key drivers of [Hb] adjustment (Fig 2, Fig 4; [38]. MCV did not vary predictably in our linear models (Fig 2D). Although increasing within-cell [Hb], or MCHC, would seem to be an alternative mechanism of increasing O_2_-carrying capacity, MCHC was less variable than other blood parameters (Table S2), likely due to physical constraints [47,48]; accordingly, MCH and MCV were strongly correlated (Table S5; Fig 4C). Contrary to recent empirical findings [30,31], adjustments to MCHC appeared to be neither substantial nor predictable with respect to elevation within or among species (Fig 2F).

**Fig 4.**
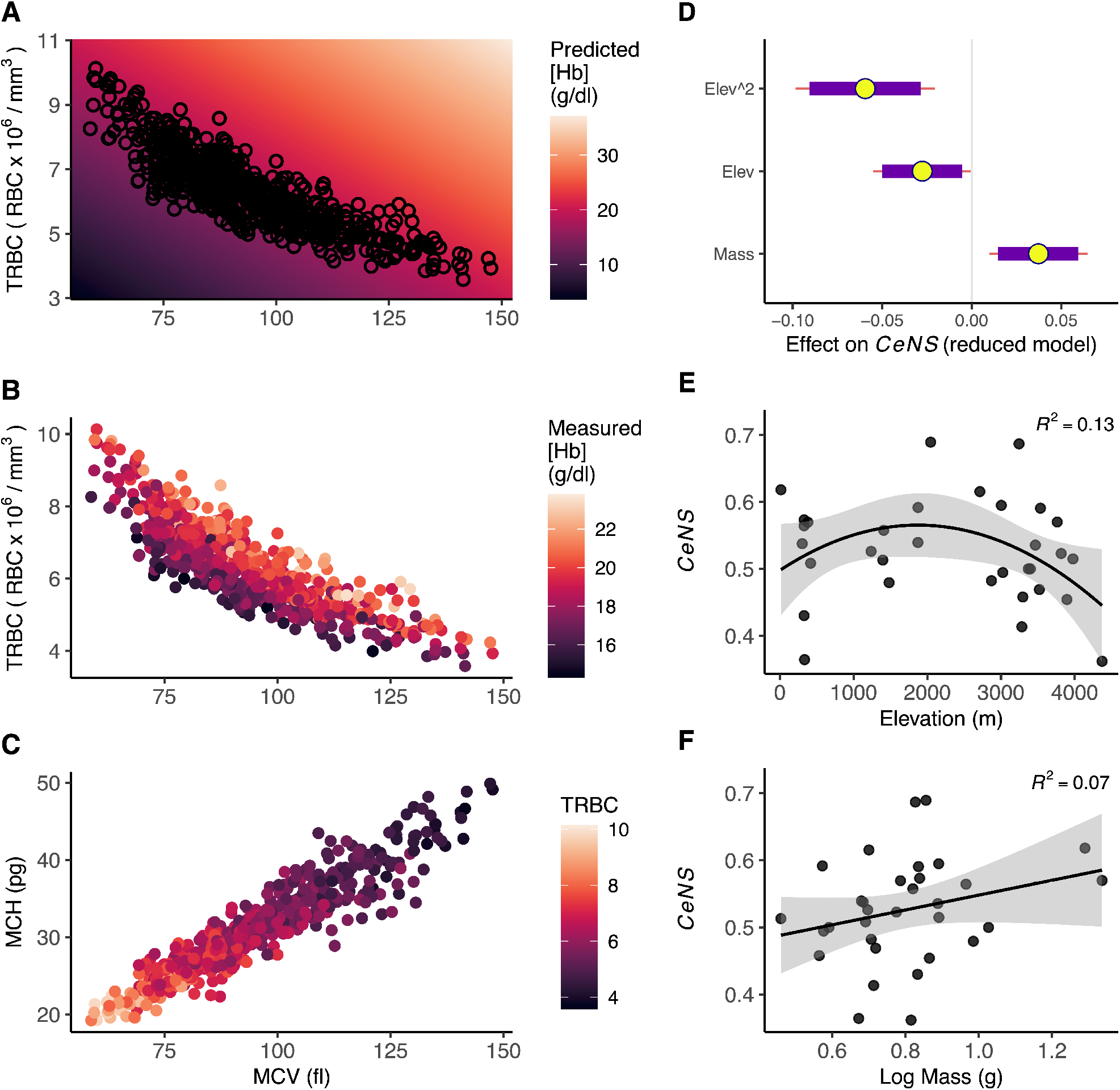
Theoretical and empirical relationships among hummingbird blood parameters underlying hemoglobin concentration ([Hb]) adjustment and predictors of the Cell Number-Size Index (*CeNS*). **(A–B)** Hummingbirds have evolved optimized blood-O_2_ carrying capacity, achieved by balancing total red blood cell count (TRBC) with mean cell volume (MCV). **(A)** Predictions for O_2_ carrying-capacity. Background is shaded by O_2_ carrying-capacity, calculated as *MCV x TRBC x 1/3 scaling factor* to represent constant MCHC. **(B)** Empirical findings. Points are colored by [Hb], with lighter colors representing higher concentrations. **(C)** Mean cell hemoglobin (MCH) and MCV are tightly linked, suggesting little variance in mean cell hemoglobin concentration (MCHC). Points are colored by TRBC (units presented on the y-axis), with lighter colors representing higher erythrocyte counts. In panels A–C, each point represents a value from a single individual. **(C)** Posterior probabilities and credible intervals from the best fitting reduced *CeNS* model. **(D)** *CeNS* shows a strong quadratic relationship with elevation, revealing that middle elevation species adjust blood-O_2_ carrying capacity differently than low- and high-elevation species. **(E)** *CeNS* increases with increasing body mass. In panels E–F, R^2^ values are reported from generalized linear models of each *x* regressed on *y*, and thus do not account for other modeled variables.

We found that MCV and MCH decreased in individuals occurring in warmer environments (Figs 3D, E), and that MCHC decreased in individuals occurring at warmer localities within a species’ range (Fig 3F). The directions of these relationships were consistent with previous experimental work: Comparisons of juvenile broiler chickens reared from ∼1-7 weeks of age in cool, ambient, and hot environments showed decreases in [Hb], Hct, MCV, and TRBC as temperature increased [49,50]. These findings add to a limited but intriguing pool of evidence that warmer ambient temperatures may lead to developmental reductions in O_2_-carrying capacity of blood.

Models that included the species grouping variable fit substantially better than those without; this effect explained 13–21% of the variance in the top model, depending on the model set (Tables S3-S4; Methods). Moderate effects of the species grouping variable suggest that variation in blood traits is partly explained by aspects of species biology that evolve too fast to have resulted in phylogenetic autocorrelation. Species-typical Hb β^A^13–β^A^83 genotype did not predict [Hb] or any other blood characteristics (Fig 3A–F), providing no evidence that evolutionary adjustments to O_2_-binding affinity affect the physical composition of blood or its variation with elevation.

### Among-species blood trait-environment variation

Using the species-level dataset, we evaluated the effects of elevation, body mass, species-typical Hb β^A^13–β^A^83 genotype, and ambient temperature and precipitation on each of our 6 blood response variables. We compared sets of models with LOOIC and WAIC (Table S3), incorporating a quadratic elevation effect for TRBC, MCV, MCH, and MCHC models based on curvature evident in the data (Fig 2G–L; Methods).

Within- and among-species findings for elevational effects were similar (Fig 3). Based on the top model in each set, for every 1,000-m increase in elevation, [Hb] is expected to increase by 0.29 g/dl, and Hct is expected to increase by 1.55 percentage points. Elevation was a consistently strong predictor of [Hb], Hct, and TRBC, all of which increased with increasing elevation (Figs 2G–I and 3A–C). The importance of the quadratic effect of elevation for MCV and MCH, and overall shape of the species-level data, suggest that O_2_ carrying capacity adjustment differs for middle-elevation species versus those found at low and high elevations (Fig 2J–K; Fig 3D–E). Neither species-typical Hb β^A^13–β^A^83 genotype, body mass, environmental temperature, nor precipitation explained variation in any species-mean blood characteristic (Fig 3).

### Mechanisms of blood-O_2_ carrying capacity adjustment

The ‘banana’ shape of the relationship between cell number versus size indicates that the highest levels of blood-O_2_ carrying capacity per unit blood are achieved at high values of TRBC or MCV, respectively. Our data support that the axis of variation in cell number versus size space is tightly constrained (Fig 4A–B), mirroring patterns evident from comparisons across multiple vertebrate classes [35,36]. Individual hummingbirds can possess few large erythrocytes or many small erythrocytes, but cell number and size must be balanced [36]. Because of the proportionately greater surface area for gas exchange, having many small cells is predicted to be advantageous for fast metabolism or when the partial pressure of O_2_ in inspired air is low [36]. Similarly, having large cells may be advantageous for modulating the time-course of gas exchange [51]. One way to shed light on this tradeoff is to test whether individuals and species, respectively, adjust O_2_ carrying capacity per unit blood primarily through increases in cell number or size.

We designed models to understand the relative contributions of cell number versus size to blood O_2_-carrying capacity per unit volume ([Hb]), within and among species over ∼22 Mya of hummingbird evolution. These models demonstrated that hummingbirds co-equally modulated cell number and cell size to adjust [Hb] (Table S6). The fact that this was true both within and among species suggests that developmental and evolutionary fine-tuning of blood-O_2_ carrying capacity occur similarly.

Using data for 32 well-sampled species, we estimated species-specific standardized TRBC and MCV coefficients (*β*) in models with [Hb] as the response variable (Fig S1), and we used these to calculate a 0–1 index that we refer to as the Cell Number-Size Index (hereafter *CeNS*). *CeNS* represents the proportional contribution of TRBC versus MCV to increasing [Hb], calculated as:

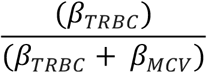

Species specific *CeNS* values ranged from 0.36-0.69 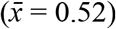 with higher values indicating a proportionally greater contribution of TRBC than MCV in adjusting [Hb]; conversely, lower values indicate a proportionally greater contribution of MCV than TRBC.

Extending our analysis, we designed a set of models to test potential predictors of *CeNS*: elevation, a quadratic elevation effect (*elevation*^*2*^; to account for notable curvature in the data), body mass, wing loading, species-typical Hb β^A^13–β^A^83 genotype, and phylogeny (see Methods). Elevation, *elevation*^*2*^, and body mass strongly predicted *CeNS* (Fig 4D-F; Table S7). We found a trend of decreasing *CeNS* with higher wing loading, but the 95% credible intervals overlapped zero (Fig S2A–B). Species-typical Hb β^A^13–β^A^83 genotype did not predict *CeNS* (Fig S3), nor did we find phylogenetic signal in *CeNS* (Fig S4).

Examining the tradeoff between cell number and volume provides a potential explanation for the striking pattern that middle elevation species tend to adjust [Hb] by different mechanisms than low and high elevation species (Fig 4E). *CeNS* was uncorrelated with TRBC coefficients (R^2^ = 0.07, *r* = -0.27) but strongly correlated with MCV coefficients (R^2^ = 0.51, *r* = -0.71), indicating that variation in *CeNS* was primarily driven by the degree to which species modulate cell size (Fig S2, Fig S5. We suggest that because low and high elevation species tend to possess relatively small cells, they may be more able to rely on shifts in cell size, rather than cell number, to increase [Hb] (Fig 2J; Fig 4). However, as cell size increased, species were more likely to approach maximum size for red blood cells, beyond which they could incur a functional detriment. As a result, species with larger red blood cells were more reliant on cell number to adjust [Hb]. For middle elevation hummingbird species that have evolved under moderate (∼15– 30%) reductions in O_2_ availability, TRBC appears to be the primary axis for adjustment of blood-O_2_ carrying capacity per unit volume of blood. Species that have evolved at the highest elevations are expected to be the best adapted to hypobaric hypoxia, and it is therefore remarkable that their mode of [Hb] adjustment (primarily by MCV) resembles that of hummingbird species in the lowlands, the putative ancestral elevation for hummingbirds [37].

### Hummingbirds at extreme elevations

Do hummingbirds at extreme high elevations follow the same rules as hummingbirds distributed at low and moderate elevations with respect to elevational blood variation? Our models encompassed the entire elevational gradient and therefore could have missed important phenomena such as threshold effects. To address this, we compared individual hummingbirds sampled above and below 4,200 m elevation, a threshold that marks the upper ∼10% of the gradient. Hummingbirds sampled from these extreme high elevations had significantly smaller cells and higher erythrocyte counts than others (Fig S6). This finding suggests that high-altitude specialist hummingbirds tend to have smaller cells, consistent with findings in other high altitude vertebrates [11,29,44,52,53]; however, this pattern was only detectable under the extreme conditions at the uppermost limits of the Andean elevational gradient. We proceeded to formally test for elevational breakpoints using segmented regression (see Methods). Evidence for objective blood trait thresholds along the elevational gradient was mixed, with breakpoint estimates for individual traits ranging from ∼1,400 to ∼4,000 m (Fig S7). Lack of consilience in these estimates suggests that trait-specific thresholds are challenging to identify and may not be generalizable across species and clades.

### Phylogenetic signal in blood traits

Phylogenetic signal was low (*λ* = 0.0; *K* range = 0.14–0.21) across all blood traits for both within- and among-species models. Blomberg’s K and Pagel’s λ results were consistent with visual assessment of continuous trait maps, even after removing species with <3 individuals sampled (Table S8; Fig S8). We additionally found no phylogenetic signal in *CeNS* (Fig S4).

Non-existent to low phylogenetic signal in blood traits within and among hummingbird species was surprising given demonstrated, phylogenetically conserved Hb genetic adaptations to elevation [21,22] and in light of strong phylogenetic signal reported from recent comparative analyses of blood traits that had broader taxonomic sampling [33,34]. However, it is likely that diverse avian clades possess different blood trait optima; in previous comparative analyses that included multiple families or orders, phylogenetic signal may have been driven primarily by differences among clades. For example, Minias et al. [33] reported that small passerines had consistently high [Hb] and Hct estimates, and that including the shorebird order, Charadriiformes, which was characterized by high [Hb] but medium Hct values, strongly influenced patterns of phylogenetic conservatism [33]. We tested for and found no interaction of clade and elevation on the six studied blood traits, with the exception of the interaction of elevation and the Hermit clade in the [Hb] model set (Fig S9). Interestingly, hermits are a large, early diverging hummingbird clade that contains diminishing species diversity with altitude (and no species that exceeds 3,000 m), and its apparent increased sensitivity of [Hb] to elevation hints at a functional explanation for this biogeographic tendency. The existence of clade-specific trait optima [33] and strong species effects, demonstrated in this study, may at first seem difficult to reconcile with our findings that there was no within-family phylogenetic signal in hummingbirds; however, we view these findings as being consistent with steady evolution of trait optima by a mean-reverting, Ornstein-Uhlenbeck process, as described by Minias et al. [33], and marked by occasional major jumps that coincide with changes in body plan or respiratory demands.

A strength of our study was that hummingbirds vary only subtly in aspects of their biology that could plausibly confound our analyses, such as metabolic rate, energy budget, torpor use, mode of locomotion, foraging strategy, or clutch size. Phylogenetic conservatism in these traits facilitates explanation of blood trait variation across the entire clade. A strong effect of species identity in spite of these intrinsic controls hints that species optimize O_2_-carrying functions of blood for subtle changes in lifestyle and environment, and on rapid timescales. This finding is consistent with the rapid diversification of human hematological responses to elevation, as illustrated by the copious variation among populations in patterns of elevational variation [19,41,42,54–56].

### Conclusions

Across ∼22 Mya of the hummingbird evolution, ambient O_2_-availability and species idiosyncrasies––not phylogenetic affinities––were the primary drivers of variation in blood traits affecting O_2_-carrying functions. In response to elevational changes in O_2_ partial pressure, hummingbirds make predictable adjustments to the physical composition of blood. Remarkably, after controlling for many potentially confounding variables, decreasing *P*O_2_ drove increases in [Hb] and Hct that were nearly identical in slope within and among hummingbird species, respectively. This implies that developmental and evolutionary optimization of blood traits to ambient *P*O_2_ occurs in fundamentally similar ways, responding to the same set of physical rules that govern gas exchange at the blood-gas barrier. In addition to O_2_ availability, species identity itself explained 13–21% of blood trait variation, suggesting that fast-evolving, unmodeled aspects of species biology underpin substantial hematological variation. Our data revealed a striking constraint space that defines the cell number-size tradeoff for hummingbird erythrocytes; within this space, individuals and species co-equally modulated cell number and size to adjust [Hb]. However, species varied in their proportional reliance on adjustments to cell number versus size (*CeNS*) to increase [Hb], and both elevation and body mass were strongly predictive of species’ tendencies. Middle-elevation species that have been evolving under moderate hypoxia adjusted [Hb] primarily by adjusting cell number; in contrast, low and high elevation species tended to adjust [Hb] by adjusting cell size, a pattern partly underpinned by hummingbirds at extreme elevations exhibiting smaller cells. Together, these results suggest that evolutionary solutions to reduced O_2_ availability can differ qualitatively in conjunction with species-specific histories of exposure to hypoxia challenges of varying severity.

## Materials and Methods

### Field Sampling

We collected data from 1,217 individuals of 77 hummingbird species from Peru and Chile from 2006–2020 (Fig 1). After filtering, our final within-species dataset included 1,151 individuals of 77 species from 75 localities. The best-sampled clades were the Coquettes (387 individuals, 20 species) and Brilliants (269 individuals, 19 species). The species with greatest sampling depth were *Metallura tyrianthina* (95 individuals), *Colibri coruscans* (73 individuals), and *Amazilia amazilia* and *Patagona gigas* (64 individuals each; Fig 1). Sampling across sexes was balanced (553 females, 592 males, and 6 unknown). Most birds sampled were adults (936 adults, 179 juveniles, and 36 unknown). Detailed data on specimens, collection localities, and blood traits are available in Witt et al. (*In prep*) and in the ARCTOS database (arctosdb.org) for reproducibility and extension of our work.

### Blood trait measurements

To calculate primary indices (hemoglobin concentration ([Hb]), hematocrit (Hct), and total red blood cell count (TRBC)), we obtained whole blood samples within ∼5 mins to ∼3 hours of capture by venipuncture on the underside of the wing and collection with heparinized microcapillary tubes and Hemocue HB201+ cuvettes. Hct (%) was measured with digital calipers after centrifuging the sealed microcapillary tube for 5 min at 13,000 r.p.m. When >1 Hct sample was taken, weighted average values were used in analyses. [Hb] (g/dL blood) was measured on ∼5 µL of blood using a HemoCue HB201+ hemoglobin photometer. The HemoCue proprietary photometric method produces values for avian blood that are ∼1.0 g/dL greater than those generated using cyanomethemoglobin spectrophotometry [57]. Accordingly, we corrected [Hb] values by subtracting 1.0 g/dL. When possible, at the time of blood collection, 10 µl whole blood was preserved in a 1:200 NaCl dilution to obtain TRBC estimates. Within 12-hours of blood collection, we pipetted a 10 µl sub-sample of blood-NaCl dilution into a hemocytometer and allowed cells to settle for 1 min. After focusing the microscope, one or more photos were obtained of cells in the hemocytometer grid. Later, cells were manually counted in the ImageJ software [58] and TRBC was estimated following standard protocols [28,59].

After drawing blood, birds were humanely killed by rapid cardiac compression. We recorded collection elevation, latitude, longitude, sex, body mass, and date. Body mass was typically recorded prior to blood sampling; however, body mass was corrected for estimated volume of blood drawn for any individuals massed after blood sampling. Each collected bird was prepared as a study skin. Specimens and tissues are housed at the Museum of Southwestern Biology (MSB) at the University of New Mexico, USA, the Centro de Ornitología y Biodiversidad (CORBIDI) in Peru, and the Universidad Pontificia Católica de Chile in Chile.

Secondary blood indices (mean cell volume (MCV), mean cell hemoglobin (MCH), and mean cell hemoglobin concentration (MCHC)) were calculated from primary indices (Table S1) and following standard protocols [28]. We examined distributions of the data using histograms and qqPlots, and we eliminated outliers that fell outside of the expected distribution and that differed from known ‘typical’ values [28,31,60]. To reduce age-related outliers, we examined values at the tails of each blood trait distribution that were associated with presence of a bursa measuring ∼2 × 2 mm. All juvenile values fell within the expected distribution of blood values. We did not age individuals by plumage, as juvenile plumage lasts much longer than juvenile blood values, which stabilize one to several months after fledging.

### Environmental and life history traits

We compiled elevational range limits for each species using a combination of published data, field guides, and specimen records from the MSB [26,61–63]. We calculated each individual’s elevational range position relative to its species elevational range breadth (hereafter ‘elevational position’) as *1 – ((maximum species elevation – individual sampling elevation)/species elevational range)*.

We incorporated species-typical Hb β^A^13–β^A^83 genotypes from published data [22,37]. For 7 species, we identified these genotypes from Hb β^A^ sequences aligned to a GenBank reference from Projecto-Garcia et al. [22]. For 12 species (*Amazilia franciae, Campylopterus villaviscensio, Eutoxeres aquila, Klais guimeti, Leucippus taczanowskii, Myrmia micrura, Phaethornis atrimentalis, Sephanoides sephaniodes, Chaetocercus mulsant, Eriocnemis vestita, Myrtis fanny*, and *Oreotrochilus leucopleurus*) species-typical Hb β^A^13–β^A^83 genotypes were inferred from sister taxa or nearest relatives. We characterized climatic variation among latitudinal and elevational gradients using 19 bioclimatic variables from the WorldClim database [64]. We downloaded rasters of each BioClim variable using the ‘raster’ package in R (version 4.0.0), cropped them to our study areas of Peru and Chile, and extracted temperature and precipitation values for each sampling locality. We used principal components analysis (PCA) for temperature (Bio1–11) and precipitation (Bio12–19) variables to create a composite measure of temperature and precipitation across the gradient. PC1 of the temperature variables (hereafter “temperature”) explained 73.4% of the variation in composite temperature, and loadings suggested that this axis corresponded to increasing temperature across sites (see GitHub). PC1 of the precipitation variables (hereafter “precipitation”) explained 79.8% of the variation in composite precipitation, and loadings suggested that this axis corresponded to increasing precipitation across sites (see GitHub). To account for multiple measurement effects (i.e., traits measured from multiple individuals within species), we calculated between- and within-species differences following Villemereuil et al. [65]. This approach uses within-group centering to separate each predictor into two components: We first calculated species mean values (i.e., between-species variability) and then subtracted the mean value of individual observations (i.e., within-species differences). These measures of within-species variation for temperature, precipitation, and body mass were included in models. Because hummingbird body masses may differ by sex, within-species differences were calculated using mean body mass of males for males, mean body mass of females for females, and species-level means for individuals of unknown sex (*n*=6).

We used wing area data from Skandalis et al. [66] to calculate wing loading in g/cm^2^, calculated as species mean body mass (g) divided by total wing area (cm^2^). We analyzed wing loading due its relevance to aerodynamic theory and hummingbird flight at high altitudes [67–70].

### Hummingbird evolutionary history

We used the rate-calibrated molecular hummingbird phylogeny from McGuire et al. [37] as a backbone to generate a phylogeny pruned to the 77 species in our dataset using the R package ‘phytools’ [71]. Two species, *Oreotrochilus leucopleurus* and *Phaethornis stuarti*, were missing from the McGuire et al. [37] tree and were added to our phylogeny using the bind.tip() function. Branch lengths and position of *O. leucopleurus* were calculated from genetic distances and recently described sister relationships [72]. Approximate branch length and position of *P. stuarti* were inferred from BirdTree.org [73,74] with divergence rate estimates from a multi-locus, time-calibrated phylogeny of Hawaiian honeycreepers [75]. We generated model-specific tree subsets by including only those tips that corresponded to the species in each modeling subset.

### Phylogenetic signal

We initially constructed phylogenetic models (results not shown), and calculated phylogenetic signal using the full and reduced models for each blood parameter, following the vignette and recommendations of P. Bürkner (https://cran.r-project.org/web/packages/brms/vignettes/brms_phylogenetics.html), using the ‘hypothesis’ method. Estimates of phylogenetic signal for all model sets (full and reduced in both within- and among species analyses) were low (λ < 0.004); these estimates are supported by visualizations of continuous trait maps of mean blood parameters (SI Appendix; Fig S8). Accordingly, we excluded phylogeny as an effect in models and evaluated phylogenetic signal using Blomberg’s K and Pagel’s λ. Results from the two tests were concordant and consistent across model subsets (Table S8).

### Within species trait-environment models

We analyzed within species variation in blood characteristics using generalized linear multilevel models in the R package brms [80]. Brms uses Markov chain Monte Carlo (MCMC) sampling to obtain draws from posterior distributions. We designed hierarchical models for each of six continuous responses variables: [Hb], Hct, TRBC, MCV, MCH, and MCHC. For each model, predictor variables included one species-level categorical variable (species-typical Hb β^A^13– β^A^83 genotype) and eight individual-level continuous variables (elevation, elevational position, body mass (log-transformed), precipitation, temperature, and within species variation in body mass, temperature, and precipitation). All variables were standardized to a mean of zero and standard deviation of one.

We compared 6 models in each model set: 1) an intercept-only null model, 2) an intercept-only model with a species grouping variable (1|species), 3) a model with all predictors and no grouping variable, 4) a model with all predictors and the species grouping variable, 5) a reduced-predictor model with no grouping variable (reduced model 3), 6) a reduced-predictor model with the species grouping variable (reduced model 4). Reduced models included only predictors whose 95% credible intervals (CIs) did not overlap zero in full models. Distribution family differed by model set: Gaussian for [Hb], Hct, and TRBC models; skew-normal for MCV and MCH models; and student’s *t*-distribution for MCHC models. We ran four Markov chains for 10,000 iterations with a burn-in period of 5,000, thinned every 10 steps.

We tested for interactions between clade and elevation by comparing three models: 1) the top within-species model for each blood parameter; 2) the top model + (1|Clade); and 3) the top model with an elev:Clade interaction. We found no meaningful interaction of elevation and clade except for elev:CladeHermit in the [Hb] model, whose 95% credible intervals did not overlap zero (Fig S9). In accordance with lack of phylogenetic signal in our data, the effect of clade explained minimal variation in model sets for each parameter (Hb=13%; Hct=8%; TRBC=4%; MCV=2%; MCH=2%; MCHC=4%).

### Among species trait-environment models

Final among-species datasets varied in the number of species they contained: ([Hb]=60 species; Hct=65 species; TRBC=58 species, MCV=60 species; MCH=59 species; MCHC=64 species). Species mean values were calculated for all species with *n*=2 or more individuals sampled. We split *Patagona gigas* into “north” and “south” haplotypes in light of evidence that the complex may be comprised of two functionally and genetically distinct populations separated by strong migratory divides [77], and we analyzed data only from individuals sampled during the breeding months to ensure correct population assignment. We designed model sets for each of our six blood traits. For each response, we analyzed distributions (qqPlots, Cook’s Distance, leverage) and removed outliers that strongly influenced fit. For each model, predictor variables included species-typical Hb β^A^13–β^A^83 genotype, mean elevation, mean body mass (log-transformed; one mean for all individuals due to balanced sampling between sexes), mean temperature, and mean precipitation. All variables were standardized to a mean of zero and standard deviation of one.

For each response, we compared: 1) an intercept-only null model, 2) a model with all predictors, and 3) a reduced-predictor model with only those predictors whose 95% CIs did not overlap zero in full models (reduced model 2). Given the notable quadratic shape of species mean data for TRBC, MCV, MCH, and MCHC (Fig 2G–L), for those model sets we additionally compared 4) a model with all predictors plus a quadratic component of elevation and 5) a reduced-predictor model with quadratic component including only those predictors whose 95% CIs did not overlap zero in the full quadratic model (reduced model 4). Distribution family differed by model set: Gaussian for [Hb; skew-normal for Hct; and student’s *t*-distribution for TRBC, MCV, MCH, and MCHC models. We ran four Markov chains for 20,000 iterations with a burn-in period of 10,000, thinned every 10 steps.

### Cell number versus size analyses

First, to assess within species variation in cell number versus size, we compared a null model (intercept only) to a single full model *[Hb] ∼ MCV + TRBC + (1*|*species)* for all individuals of 59 species that had [Hb], MCV, and TRBC measurements, distinguishing between *Patagona gigas* “north” and “south”. The species grouping variable allowed us to account for repeated sampling of individuals within a species and to hold species identity constant. Second, to evaluate among-species variation in cell number versus size, we generated a mean dataset for each species with ≥3 individuals sampled (48 species total). We again compared a null (intercept only) model to a single full model *[Hb] ∼ MCV + TRBC*. Third, to assess how individual species differ in their relative contributions of MCV and TRBC to increasing [Hb], we repeated our approach for individual species: For each species with ≥8 sample individuals (32 species total), we generated a species-specific data subset, carefully and conservatively evaluated influential outlier points, and ran a single model: *[Hb] ∼ MCV + TRBC* (Fig S1). For all three model sets (within species, among species, and single species), both predictors were standardized to a mean of zero and standard deviation of one.

Using standardized, model-estimated coefficients from each single-species model, for each species we calculated the Cell Number-Size Index (hereafter *CeNS*) as described in the main text. *CeNS* ranged from 0.0–1.0 and represented the proportional contribution of TRBC relative to MCV in driving [Hb], with higher values corresponding to greater relative contributions of cell number versus cell size.

We then analyzed the factors that contribute to variation in *CeNS*. Our model response variable was *CeNS* (0–1 index). Predictors, all species mean values, included elevation, a quadratic elevation term “elevation^2^” to account for notable curvature in the data, body mass (log-transformed), wing loading, and species-typical Hb β^A^13–β^A^83 genotype. We compared six models: 1) an intercept-only null model, 2) a model with core predictors (elevation, body mass, and wing loading), 3) a model with core predictors plus elevation^2^, 4) a model with core predictors plus elevation^2^ and species-typical Hb β^A^13–β^A^83 genotype, 5) a reduced version of model 3, and 6) a reduced version of model 4. Reduced models included only those predictors whose 95% CIs did not overlap zero in full models. In addition to model fit diagnostics, we assessed correlations between variables by analyzing correlation matrices and by regressing *CeNS* on each R^2^ values from standardized species-specific *CeNS* models, unstandardized TRBC and MCV beta estimates, and raw TRBC and MCV values (Fig S5). Distribution family differed by model set: Gaussian for individual-level, species mean, and *CeNS* prediction models, and student’s *t*-distribution for species-specific models. For all *CeNS* models we ran four Markov chains for 20,000 iterations with a burn-in period of 10,000, thinned every 10 steps. We assessed phylogenetic signal with Blomberg’s K, Pagel’s λ, and visually with contmaps (Fig S4).

### Model comparison and diagnostics

All models and sets were run using the brm() function with default priors (weakly informative priors for the intercept and sigma; improper flat priors for predictors). To assess convergence we visually examined trace plots, checked diagnostics, and confirmed that Rhat was <1.01 [76]. We used a posterior predictive check to assess whether model assumptions yielded good approximations of the data generating process. There were no problems in any of our models resulting from Bayesian fraction of missing information or divergent transitions. We compared models and verified fits using the widely applicable information criterion (WAIC) [78], Bayesian expected log pointwise predictive densities (ELPD) [79], and approximate leave-one-out cross-validation information criterion based on the posterior likelihoods (LOOIC; Tables S3, S6). We estimated fixed effects (means and 95% CIs) from the posterior distributions for each predictor. For relevant models, we estimated the proportion of the total variance attributed to the species grouping variable, which accounts for unique aspects of blood parameters that are not captured by the individual, life history, or environmental predictors.

### Breakpoint analysis

We tested for objective thresholds in each blood trait using piecewise linear regression of elevation (standardized) regressed on each blood trait. In this procedure, we first fit a simple linear regression model, iteratively searched for objective “breakpoints” along our sampled elevational gradient, selected the breakpoint with the lowest residual error, and refit the linear regression model with separate estimates for slopes and intercepts below and above the breakpoint, respectively. We constrained the model to estimate breakpoints that were each 400 m from the lowest and highest points in our data (i.e., from 400 m to 4,178 m in elevation), to avoid biasing breakpoints towards small sample sizes and each low and high elevations, and anomalous species at both ends of the distribution. We evaluated goodness of fit using fit metrics and by visually assessing QQplots, Cook’s D, leverage, and residual distributions. We compared models using AIC values. All breakpoint models fit significantly better than simple linear regression models except for the TRBC breakpoint model; this model did fit better in all respects than the simple linear regression model, but the fit was not significant at the *p*=0.05 level. We used a one-way ANOVA to compare whether individuals below and above model-estimated breakpoints differed in their blood characteristics (Fig S7).

## Data and code availability

All data and code for analyses are available in an open GitHub repository: https://github.com/jlwilliamson/ComparativeHummingbirdBlood. Specimen data are available from the Arctos database (https://www.arctosdb.org). Additional detailed data on specimens, collection localities, and blood values are reported in a forthcoming data paper, Witt et al. (*In Prep*).

## Acknowledgements

We thank Luis Alza, Christopher Barger, Selina Bauernfeind, Matthew Baumann, Elizabeth Beckman, Phred Benham, José Miguel Bogdanovich, Francisco Bozinovic, Homan Castillo Benitez, Jessica Castillo, Marlon Chagua, Jennifer Clark, Mariela Combe, Lida Crooks, Robert Driver, Shane DuBay, Margarita Espinoza, L. Monica Flores, Chauncey Gadek, Ariel Gaffney, Spencer Galen, Avia González-Méndez, Paige Handley, Zachary Hanna, Mike Hartshorne, José Ernesto Huaroto Tornero, Andrew Johnson, Matthew Jones, Daniel Lane, LyAndra Lujan, Mattías Marzfeld, Sabrina McNew, Kyana Montoya, Jano Nuñez-Zapata, Iris Olivas, Paloma Ordoñez, Alessandra Quiñonez, Javier Reinoso, Natalia Ricote, C. Gregory Schmitt, Donna Schmitt, C. Jonathan Schmitt, Frank Solano Bravo, Dora Susanibar, William Talbot, Jorge Tiravanti C., Abraham Urbay T., Thomas Valqui, Walter Vargas Campos, Karen Verde-Guerra, Natalie Wright, and Alfredo Zelada. We are grateful to the following communities in Peru for site access: San Pedro de Casta, Comunidad Campesina Santiago de Carampoma, Ocros, Macate, Oncoy, Yanahara, Huacarpay, Sianbal, Plataforma, Agua Azul, Incahuasi, and Anchicha. This research was funded by the National Science Foundation (DEB-0543556, DEB-1146491, and DBI-1907353), and dissertation research grants to JLW from the Nuttall Ornithological Club Blake-Nuttall Fund Grants (2016 & 2018), the American Philosophical Society Lewis & Clark Grant, the Explorers Club Exploration Fund Grant, American Museum of Natural History Chapman grants (2017 & 2019), the UNM Biology Graduate Student Association Graduate Research Allocations Committee grant, UNM Graduate and Professional Student Association grants (2017–2021), the UNM Latin American & Iberian Institute Field Research Grant, UNM Department of Biology Grove Research Scholarships (2018, 2019, and 2021), and the UNM Department of Biology Melinda Bealmear Memorial Scholarship and Dr. Jones & Dr. Wong Scholarship. Permits in Chile were granted by Servicio Agrícola y Ganadero (SAG; RE Noºs: 7593/2016, 6817/2017, 7903/2018, 6691/2018, 6692/2018, 6693/2018, 6694/2018). Permits in Peru were granted by Servicio Nacional Forestal y de Fauna Silvestre (SERFOR; RDG Noºs: 004-2007-INRENA-IFFS-DCB, 135-2009-AG-DGFFS-DGEFFS, 0377-2010-AG-DGFFS-DGEFFS, 0199-2012-AG-DGFFS-DGEFFS, and 006-2013-MINAGRI-DGFFS/DGEFFS, 244-2020-MINAGRI-SERFOR/DGGSPFFS-DGSPFS). UNM IACUC approval was granted under protocols 16-200596-MC and 19-200804-MC.

## Author Contributions

JLW, EBL, EB, JAM, RD, and CCW designed the research and methods; JLW, EB, AS, JAM, RD, and CCW collected the data; JLW analyzed the data; JLW, JAM, RD, and CCW acquired funds and resources; and JLW and CCW wrote the paper with input from all authors.

## References

1. McDowall RM. Variation in vertebral number in galaxiid fishes (Teleostei: Galaxiidae): A legacy of life history, latitude and length. Environ Biol Fishes. 2003;66: 361–381. doi:10.1023/A:1023902922843

2. Jordan DS. Relations of temperature to vertebrae among fishes. Proc United States Natl Museum. 1891;14: 107–120. doi:10.5479/si.00963801.14-845.107

3. Blackburn TM, Gaston KJ, Loder N. Geographic gradients in body size: A clarification of Bergmann’s rule. Divers Distrib. 1999;5: 165–174. doi:10.1046/j.1472-4642.1999.00046.x

4. De Queiroz A, Ashton KG. The phylogeny of a species-level tendency: Species heritability and possible deep origins of Bergmann’s rule in tetrapods. Evolution (N Y). 2004;58: 1674–1684. doi:10.1111/j.0014-3820.2004.tb00453.x

5. McGlothlin JW, Kobiela ME, Wright H V., Mahler DL, Kolbe JJ, Losos JB, et al. Adaptive radiation along a deeply conserved genetic line of least resistance in Anolis lizards. Evol Lett. 2018;2: 310–322. doi:10.1002/evl3.72

6. Agrawal AA. A scale-dependent framework for trade-offs, syndromes, and specialization in organismal biology. Ecology. 2020;101: 1–24. doi:10.1002/ecy.2924

7. Züst T, Rasmann S, Agrawal AA. Growth-defense tradeoffs for two major anti-herbivore traits of the common milkweed Asclepias syriaca. Oikos. 2015;124: 1404–1415. doi:10.1111/oik.02075

8. Agrawal AA, Fishbein M. Phylogenetic escalation and decline of plant defense strategies. Proc Natl Acad Sci U S A. 2008;105: 10057–10060. doi:10.1073/pnas.0802368105

9. Enquist BJ, Norberg J, Bonser SP, Violle C, Webb CT, Henderson A, et al. Scaling from Traits to Ecosystems: Developing a General Trait Driver Theory via Integrating Trait-Based and Metabolic Scaling Theories. In: Samraat Pawar, Guy Woodward AID, editor. Advances in Ecological Research. Elsevier Ltd.; 2015. pp. 249–318. doi:10.1016/bs.aecr.2015.02.001

10. Borras A, Cabrera J, Senar JC. Hematocrit Variation in Response to Altitude Changes in Wild Birds: A Repeated-Measures Design. Condor. 2010;112: 622–626. doi:10.1525/cond.2010.090113

11. Tufts DM, Revsbech IG, Cheviron ZA, Weber RE, Fago A, Storz JF. Phenotypic plasticity in blood-oxygen transport in highland and lowland deer mice. J Exp Biol. 2013;216: 1167–1173. doi:10.1242/jeb.079848

12. Levett D, Radford E, Menassa D, Graber E, Morash A, Hoppeler H, et al. Acclimatization of skeletal muscle mitochondria to high-altitude hypoxia during an ascent of Everest. FASEB J. 2012;26: 1431–1441. doi:10.1096/fj.11-197772

13. Storz JF, Scott GR, Cheviron ZA. Phenotypic plasticity and genetic adaptation to high-altitude hypoxia in vertebrates. J Exp Biol. 2010;213: 4125–36. doi:10.1242/jeb.048181

14. Pittman RN. Chapter 4: Oxygen Transport. Regulation of Tissue Oxygentation. Morgan & Claypool Life Sciences; 2011. Available: https://www.ncbi.nlm.nih.gov/books/NBK54103/

15. Stembridge M, Williams AM, Gasho C, Dawkins TG, Drane A, Villafuerte FC, et al. The overlooked significance of plasma volume for successful adaptation to high altitude in Sherpa and Andean natives. Proc Natl Acad Sci U S A. 2019;116: 16177–16179. doi:10.1073/pnas.1909002116

16. Storz JF. Genes for High Altitudes. Science (80-). 2010;329: 40–41. doi:10.1126/science.1192481

17. Heinicke K, Prommer N, Cajigal J, Viola T, Behn C, Schmidt W. Long-term exposure to intermittent hypoxia results in increased hemoglobin mass, reduced plasma volume, and elevated erythropoietin plasma levels in man. Eur J Appl Physiol. 2003;88: 535–543. doi:10.1007/s00421-002-0732-z

18. Scott GR, Milsom WK. Flying high: A theoretical analysis of the factors limiting exercise performance in birds at altitude. Respir Physiol Neurobiol. 2006;154: 284–301. doi:10.1016/j.resp.2006.02.012

19. Beall CM. Two routes to functional adaptation: Tibetan and Andean high-altitude natives. Proc Natl Acad Sci. 2007;104: 8655–8660. doi:10.1073/pnas.0701985104

20. Simonson TS, Yang Y, Huff CD, Yun H, Qin G, Witherspoon DJ, et al. Genetic Evidence for High-Altitude Adaptation in Tibet. Science (80-). 2010;329: 72–75. doi:10.1126/science.1189406

21. Natarajan C, Hoffmann FG, Weber RE, Fago A, Witt CC, Storz JF. Predictable convergence in hemoglobin function has unpredictable molecular underpinnings. Science (80-). 2016;354: 336–339.

22. Projecto-Garcia J, Natarajan C, Moriyama H, Weber RE, Fago A, Cheviron ZA, et al. Repeated elevational transitions in hemoglobin function during the evolution of Andean hummingbirds. Proc Natl Acad Sci. 2013;110: 20669–20674.

23. Natarajan C, Projecto-Garcia J, Moriyama H, Weber RE, Muñoz-Fuentes V, Green AJ, et al. Convergent Evolution of Hemoglobin Function in High-Altitude Andean Waterfowl Involves Limited Parallelism at the Molecular Sequence Level. PLoS Genet. 2015;11: 1–25. doi:10.1371/journal.pgen.1005681

24. Storz JF, Moriyama H. Mechanisms of hemoglobin adaptation to high altitude hypoxia. High Alt Med Biol. 2008;9: 148–157.

25. Zhu X, Guan Y, Qu Y, David G, Song G, Lei F. Elevational divergence in the great tit complex revealed by major hemoglobin genes. Curr Zool. 2017;64: 455–464. doi:10.1093/cz/zox042

26. Williamson JL, Witt CC. Elevational niche-shift migration: Why the degree of elevational change matters for the ecology, evolution, and physiology of migratory birds. Ornithology. 2021;138: 1–26. doi:10.1093/ornithology/ukaa087

27. Storz JF. Hemoglobin–oxygen affinity in high-altitude vertebrates: is there evidence for an adaptive trend? J Exp Biol. 2016;219: 3190–3203. doi:10.1242/jeb.127134

28. Campbell TW, Ellis CK. Avian and Exotic Animal Hematology and Cytology. Blackwell Publishing Inc.; 2007. doi:10.1017/CBO9781107415324.004

29. Zhang H, Wu CX, Chamba Y, Ling Y. Blood characteristics for high altitude adaptation in Tibetan chickens. Poult Sci. 2007;86: 1384–1389. doi:10.1093/ps/86.7.1384

30. Barve S, Dhondt AA, Mathur VB, Cheviron ZA. Life-history characteristics influence physiological strategies to cope with hypoxia in Himalayan birds. Proc R Soc B. 2016;283: 1–8. doi:dx.doi.org/10.1098/rspb.2016.2201

31. Dubay SG, Witt CC. Differential high-altitude adaptation and restricted gene flow across a mid-elevation hybrid zone in Andean tit-tyrant flycatchers. Mol Ecol. 2014;23: 3551–3565. doi:10.1111/mec.12836

32. Linck EB, Williamson JL, Bautista E, Beckman EJ, Benham PM, DuBay SG, et al. Blood variation implicates respiratory limits on elevational ranges of Andean birds. bioRxiv. 2021; 1–28. doi:https://doi.org/10.1101/2021.09.30.462673

33. Minias P. Ecology and evolution of blood oxygen-carrying capacity in birds. Am Nat. 2020;195: 788–801. doi:10.1086/707720

34. Yap KN, Tsai OH-I, Williams TD. Haematological traits co-vary with migratory status, altitude and energy expenditure: a phylogenetic, comparative analysis. Sci Rep. 2019;9: 6351. doi:10.1038/s41598-019-42921-4

35. Wintrobe MM. Variation in the size and hemoglobin content of erythrocytes in the blood of various vertebrates. Folia Haematol. 1933;51: 32–59.

36. Hawkey CM, Bennett PM, Gascoyne SC, Hart MG, Kirkwood JK. Erythrocyte size, number and haemoglobin content in vertebrates. Br J Haematol. 1991;77: 392–397. doi:10.1111/j.1365-2141.1991.tb08590.x

37. McGuire JA, Witt CC, Remsen J V, Corl A, Rabosky DL, Altshuler DL, et al. Molecular phylogenetics and the diversification of hummingbirds. Curr Biol. 2014;24: 1–7. doi:10.1016/j.cub.2014.03.016

38. Suarez RK. Hummingbird flight: Sustaining the highest mass-specific metabolic rates among vertebrates. Experientia. 1992;48: 565–570. doi:10.1007/BF01920240

39. Graham CH, Parra JL, Rahbek C, McGuire JA. Phylogenetic structure in tropical hummingbird communities. Proc Natl Acad Sci. 2009;106: 19673–19678. doi:10.1073/pnas.0912879107

40. Buermann W, Chaves JA, Dudley R, Mcguire JA, Smith TB, Altshuler DL. Projected changes in elevational distribution and flight performance of montane Neotropical hummingbirds in response to climate change. Glob Chang Biol. 2011;17: 1671–1680. doi:10.1111/j.1365-2486.2010.02330.x

41. Beall CM, Brittenham GM, Strohl KP, Blangero J, Williams-Blangero S, Goldstein MC, et al. Hemoglobin concentration of high-altitude Tibetans and Bolivian Aymara. Am J Phys Anthropol. 1998;106: 385–400.

42. Beall CM. Andean, Tibetan, and Ethiopian patterns of adaptation to high-altitude hypoxia. Integr Comp Biol. 2006;46: 18–24. doi:10.1093/icb/icj004

43. Hammond KA, Szewczak JOE, Król E. Effects of altitude and temperature on organ phenotypic plasticity along an altitudinal gradient. J Exp Biol. 2001;204: 1991–2000.

44. Lu S, Xin Y, Tang X, Yue F, Wang H, Bai Y, et al. Differences in hematological traits between high-and low-altitude lizards (Genus Phrynocephalus). PLoS One. 2015;10: 1–16. doi:10.1371/journal.pone.0125751

45. Hadley NF, Burns TA. Intraspecific Comparison of the Blood Properties of the Side-Blotched Lizard, Uta stansburiana. Copeia. 1968;1968: 737–740.

46. Dawson NJ, Alza L, Nandal G, Scott GR, McCracken KG. Convergent changes in muscle metabolism depend on duration of high-altitude ancestry across andean waterfowl. Elife. 2020;9: 1–35. doi:10.7554/eLife.56259

47. Fischer SL, Fischer SP. Mean Corpuscular Volume. JAMA Intern Med. 1983;143: 282–283. doi:10.1001/archpedi.1981.02130320078028

48. Rose MS. Epitaph for the M.C.H.C. Br Med J. 1971;4: 169.

49. Donkoh A. Ambient temperature: a factor affecting performance and physiological response of broiler chickens. Int J Biometeorol. 1989;33: 259–265. doi:10.1007/BF01051087

50. Moye RJ, Washburn KW, Huston TM. Effects of environmental temperature on erythrocyte numbers and size. Poult Sci. 1969;48: 1683–1686. doi:10.3382/ps.0481683

51. Glomski CA, Pica A. The Avian Erythrocyte: Its Phylogenetic Odyssey. 1st ed. Enfield, New Hampshire, USA: Science Publishers; 2011.

52. Ruiz G, Rosenmann M, Veloso A. Altitudinal distribution and blood values in the toad, Bufo spinulosus Wiegmann. Comp Biochem Physiol -- Part A Physiol. 1989;94: 643–646. doi:10.1016/0300-9629(89)90609-9

53. Yamaguchi K, Jürgens KD, Bartels H, Piiper J. Oxygen transfer properties and dimensions of red blood cells in high-altitude camelids, dromedary camel and goat. J Comp Physiol B Biochem Syst Environ Physiol. 1987;157: 1–9. doi:10.1007/BF00702722

54. Brutsaert TD, Araoz M, Soria R, Spielvogel H, Haas JD. Higher arterial oxygen saturation during submaximal exercise in Bolivian Aymara compared to European sojourners and Europeans born and raised at high altitude. Am J Phys Anthropol. 2000;113: 169–181. doi:10.1002/1096-8644(200010)113:2<169::AID-AJPA3>3.0.CO;2-9

55. Grimminger J, Richter M, Tello K, Sommer N, Gall H, Ghofrani HA. Thin Air Resulting in High Pressure: Mountain Sickness and Hypoxia-Induced Pulmonary Hypertension. Can Respir J. 2017; 1–17. doi:10.1155/2017/8381653

56. Guo Y-B, He Y-X, Cui C-Y, Ouzhuluobu, Baimakangzhuo, Duojizhuoma, et al. GCH1 plays a role in the high-altitude adaptation of Tibetans. Zool Res. 2017;38: 155–162. doi:10.24272/j.issn.2095-8137.2017.037

57. Simmons P, Lill A. Development of parameters influencing blood oxygen carrying capacity in the welcome swallow and fairy martin. Comp Biochem Physiol - A Mol Integr Physiol. 2006;143: 459–468. doi:10.1016/j.cbpa.2005.12.018

58. Rasband WS. ImageJ. Maryland, USA: U.S. National Institutes of Health; Available: https://imagej.nih.gov/ij/

59. Samour J. Diagnostic Value of Hematology. In: Harrison G, Lightfoot T, editors. Clinical Avian Medicine, Volume II. 2005. pp. 587–610. Available: http://avianmedicine.net/publication_cat/clinical-avian-medicine/

60. Elarabany N. A comparative study of some haematological and biochemical parameters between two species from the Anatidae family within migration season. J Basic Appl Zool. 2018;79. doi:10.1186/s41936-018-0044-4

61. Parker TAI, Stotz DF, Fitzpatrick JW. Ecological and Distributional Databases. In: D.F. Stotz, J.W. Fitzpatrick, T.A. Parker, III Dkm, editor. Neotropical Birds: Ecology and Conservation. Chicago, IL, USA: University of Chicago Press; 1996.

62. Jaramillo A. Birds of Chile. Princeton, New Jersey, USA: Princeton University Press; 2003.

63. Schulenberg T, Thomas T, Stotz D. Birds of Peru: revised and updated edition. Princeton Univeristy Press; 2010.

64. Hijmans RJ, Cameron SE, Parra JL, Jones PG, Jarvis A. Very high resolution interpolated climate surfaces for global land areas. Int J Climatol. 2005;25: 1965–1978. doi:10.1002/joc.1276

65. Villemereuil P de, Nakagawa S. Modern phylogenetic comparative methods and their application in evolutionary biology. Mod Phylogenetic Comp Methods their Appl Evol Biol. 2014; 1–552. doi:10.1007/978-3-662-43550-2

66. Skandalis DA, Segre PS, Bahlman JW, Groom DJE, Welch KC, Witt CC, et al. The biomechanical origin of extreme wing allometry in hummingbirds. Nat Commun. 2017;8: 1–9. doi:10.1038/s41467-017-01223-x

67. Dudley R. Mechanisms and Implications of Animal Flight Maneuverability. Integr Comp Biol. 2002;42: 135–140. doi:10.1093/icb/42.1.135

68. Roslyn Dakin, Paolo S. Segre, Andrew D. Straw DLA. Morphology, muscle capacity, skill, and maneuvering ability in hummingbirds. Science (80-). 2018;657: 653–657.

69. Lee SY, Scott GR, Milsom WK. Have wing morphology or flight kinematics evolved for extreme high altitude migration in the bar-headed goose? Comp Biochem Physiol Part C. 2008;148: 324–331. doi:10.1016/j.cbpc.2008.05.009

70. Altshuler DL, Dudley R, McGuire JA. Resolution of a paradox: Hummingbird flight at high elevation does not come without a cost. Proc Natl Acad Sci U S A. 2004;101: 17731–17736. doi:10.1073/pnas.0405260101

71. Revell LJ. phytools: an R package for phylogenetic comparative biology (and other things). Methods Ecol Evol. 2012;3: 217–223. doi:10.1111/j.2041-210X.2011.00169.x

72. Sornoza-Molina F, Freile JF, Nilsson J, Krabbe N, Bonaccorso E. A striking, critically endangered, new species of hillstar (Trochilidae: Oreotrochilus) from the southwestern Andes of Ecuador. Auk. 2018;135: 1146–1171. doi:10.1642/AUK-18-58.1

73. Rubolini D, Liker A, Garamszegi LZ, Moller AP, Saino N. Using the BirdTree. org website to obtain robust phylogenies for avian comparative studies: A primer. Curr Zool. 2015;61: 959–965.

74. Jetz W, Thomas GH, Joy JB, Hartmann K, Mooers AO. The global diversity of birds in space and time. Nature. 2012;491: 444–498. doi:10.1038/nature11631

75. Lerner HRL, Meyer M, James HF, Hofreiter M, Fleischer RC. Multilocus resolution of phylogeny and timescale in the extant adaptive radiation of Hawaiian honeycreepers. Curr Biol. 2011;21: 1838–1844. doi:10.1016/j.cub.2011.09.039

76. Bürkner PC. brms: An R package for Bayesian multilevel models using Stan. J Stat Softw. 2017;80. doi:10.18637/jss.v080.i01

77. Williamson JL, Witt CC. A lightweight backpack harness for tracking hummingbirds. J Avian Biol. 2021; 1–9. doi:10.1111/jav.02802

78. Watanabe S. Asymptotic equivalence of Bayes cross validation and widely applicable information criterion in singular learning theory. J Mach Learn Res. 2010;11: 3571–3594.

79. Vehtari A, Gelman A, Gabry J. Practical Bayesian model evaluation using leave-one-out cross-validation and WAIC. Stat Comput. 2017; 1413–1432.

