## Supplementary Information for "Hummingbird blood traits track oxygen availability across space and time"

### Supplementary Methods

*Within and among species models*

We used LOOIC and WAIC to compare models in both within and among species model sets (Table S3). For within species models, the full predictor models with species grouping variable had the lowest (best) WAIC and ELPD values for [Hb] ($R^2$ = 0.21), Hct ($R^2$ = 0.17), and TRBC ($R^2$ = 0.17) models. For MCV, the reduced model with temperature and the species grouping variable ranked the highest ($R^2$ = 0.094) and for MCHC, the reduced model that included within species variation in both body mass and temperature plus the species grouping variable ranked the highest ($R^2$ = 0.12). Both top MCV and MCHC models ranked similarly to the full model with species grouping variable, as well as the null model. For MCH, the reduced model with elevational position, elevation, and the species grouping variable ranked the highest and was comparable to the null model ($R^2$ = 0.092; Table S3).

For among species models the [Hb] model with the best (lowest) WAIC and ELPD values was a reduced model with only elevation ($R^2$ = 0.18), though this model was only marginally better than the full model with all predictors ($R^2$ = 0.33; Table S3). The top Hct model was the full model with all predictors ($R^2$ = 0.24; Table S3). For MCH, the top model was a reduced model containing elevation and a quadratic parameter for elevation, both of which were important predictors ($R^2$ = 0.12; Tables S3-S4; Figures 2K and 3E). For TRBC, MCV, and

MCHC model sets, the intercept-only model received the strongest support, though other models were closely ranked (Table S5).

*Cell Size versus Number Index (CeNS) models*

For both within- and among-species Cell Number versus Size Index (*CeNS*) models, full models fit substantially better than the intercept-only null model (full within species model $R^2$ = 0.56; full among species model $R^2$ = 0.62; Table S6). Based on model estimates, for every increase in TRBC of one million cells per µl, we expect that [Hb] will increase by 1.81 g/dl within species and 2.15 g/dl among species, and for every increase in MCV of one fl, we expect that [Hb] will increase by 0.12 g/dl within species and 0.14 g/dl among species.

In our analysis to assess the factors that contribute to variation in *CeNS*, we compared six models using LOOIC and WAIC (Table S7). The model with elevation, $elevation^2$, body mass, and wing loading received the strongest support ($R^2$ = 0.38) but was closely ranked to a model including only elevation, $elevation^2$, and body mass ($R^2$ = 0.33; Figure 4D; Table S7). Models with $elevation^2$ fit substantially better than those without. All but *Coeligena coeligena* showed a positive relationship between *CeNS* and predictors (Figure S1).

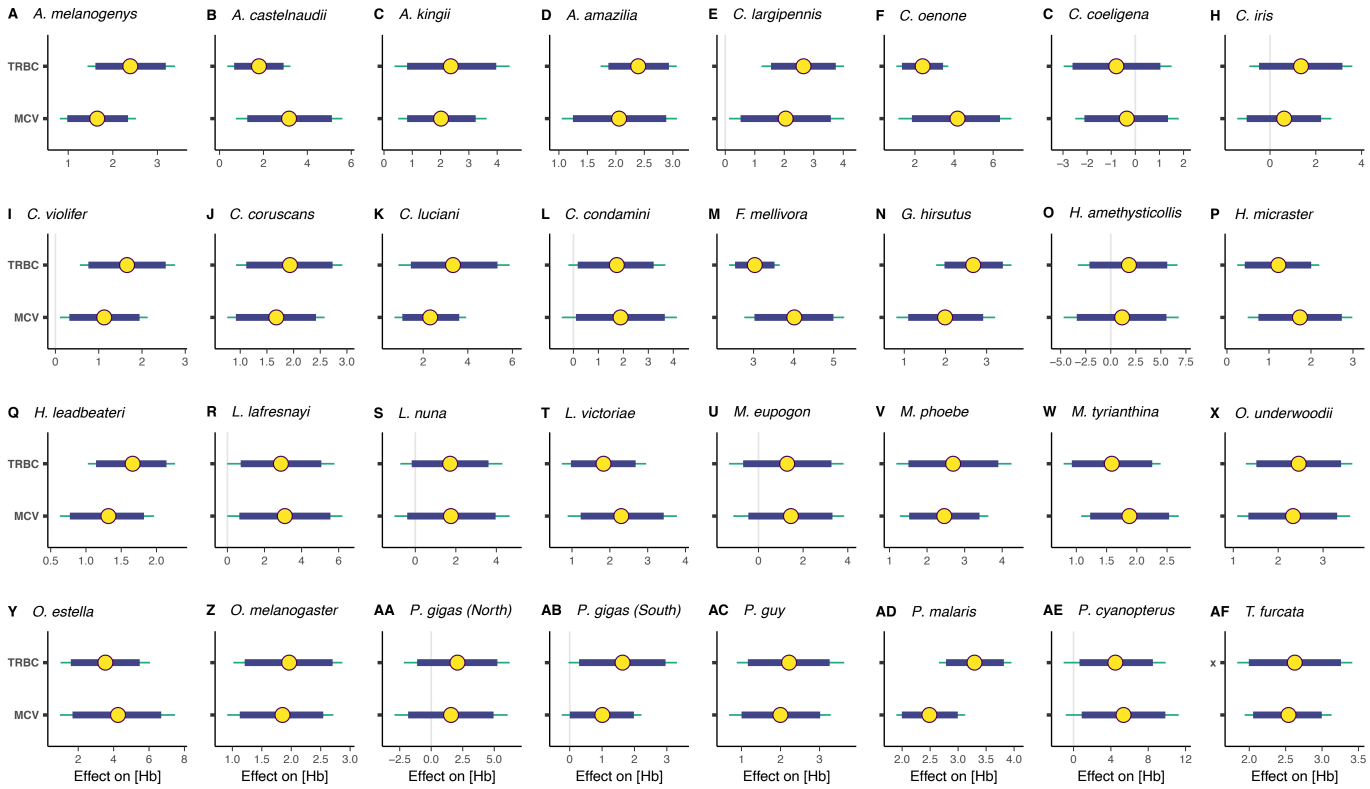


**Figure S1. (A–AF)** Posterior probabilities and credible intervals from species-specific cell size versus number models, the coefficients from which were used to calculate the Cell Size-Number Index (*CeNS*). Data from a single species are presented in each panel. Circles indicate parameter estimates ($\beta$s), and thick and thin bars illustrate 89% and 95% credible intervals, respectively.

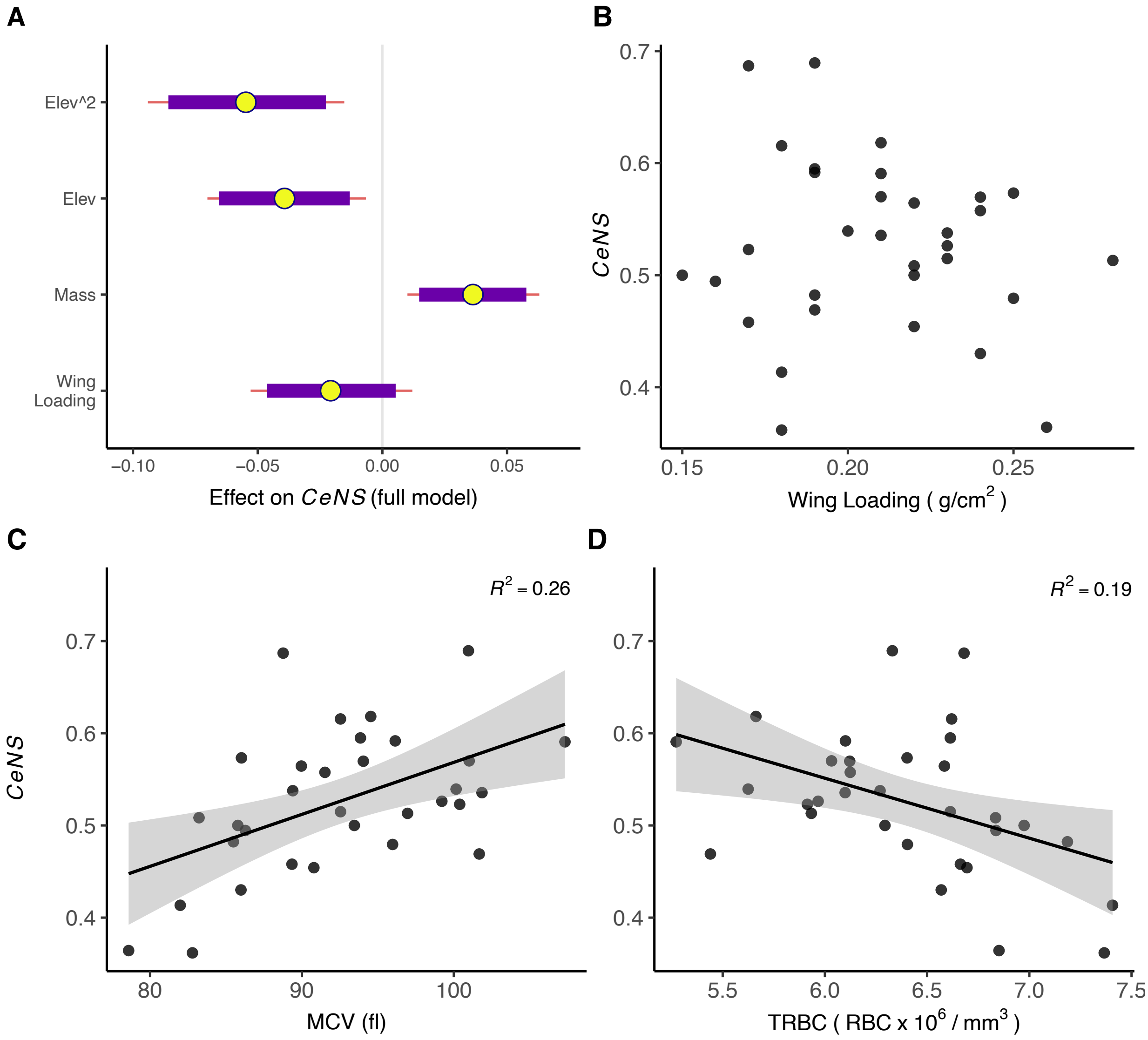


**Figure S2. (A)** Posterior probabilities and credible intervals from the best fitting full *CeNS* model (reduced model shown in Figure 4). **F)** *CeNS* shows a marginal decrease with increased wing loading. **(C–D)** Effects of mean cell volume (MCV) and total red blood cell count (TRBC) on *CeNS*, elucidating how, once cells reach a certain size, TRBC becomes the primary mechanism through which birds adjust [Hb]. $R^2$ values are reported from generalized linear models of each *x* regressed on *y*, and thus do not account for other modeled variables (relevant only to panel B).

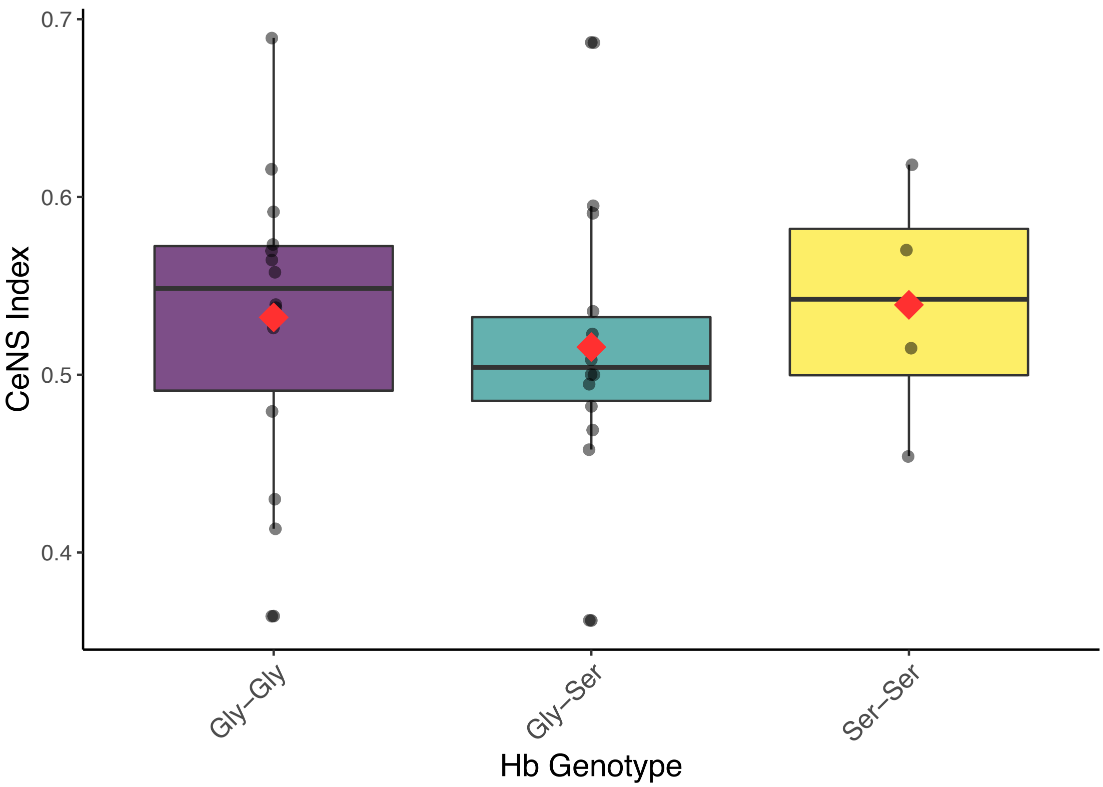


**Figure S3.** Hemoglobin $\beta^{A}$-13$_{GLY\rightarrow SER}$ and Hb $\beta^{A}$-83 $_{GLY\rightarrow SER}$ genotype (or "Hb genotype") did not predict *CeNS*. *CeNS* ranged from 0 to 1, with higher values indicating a proportionally greater contribution of TRBC than MCV in increasing [Hb]; similarly, lower values indicate proportionally greater contributions of MCV than TRBC in increasing [Hb]. Each point is a mean value for a single species.

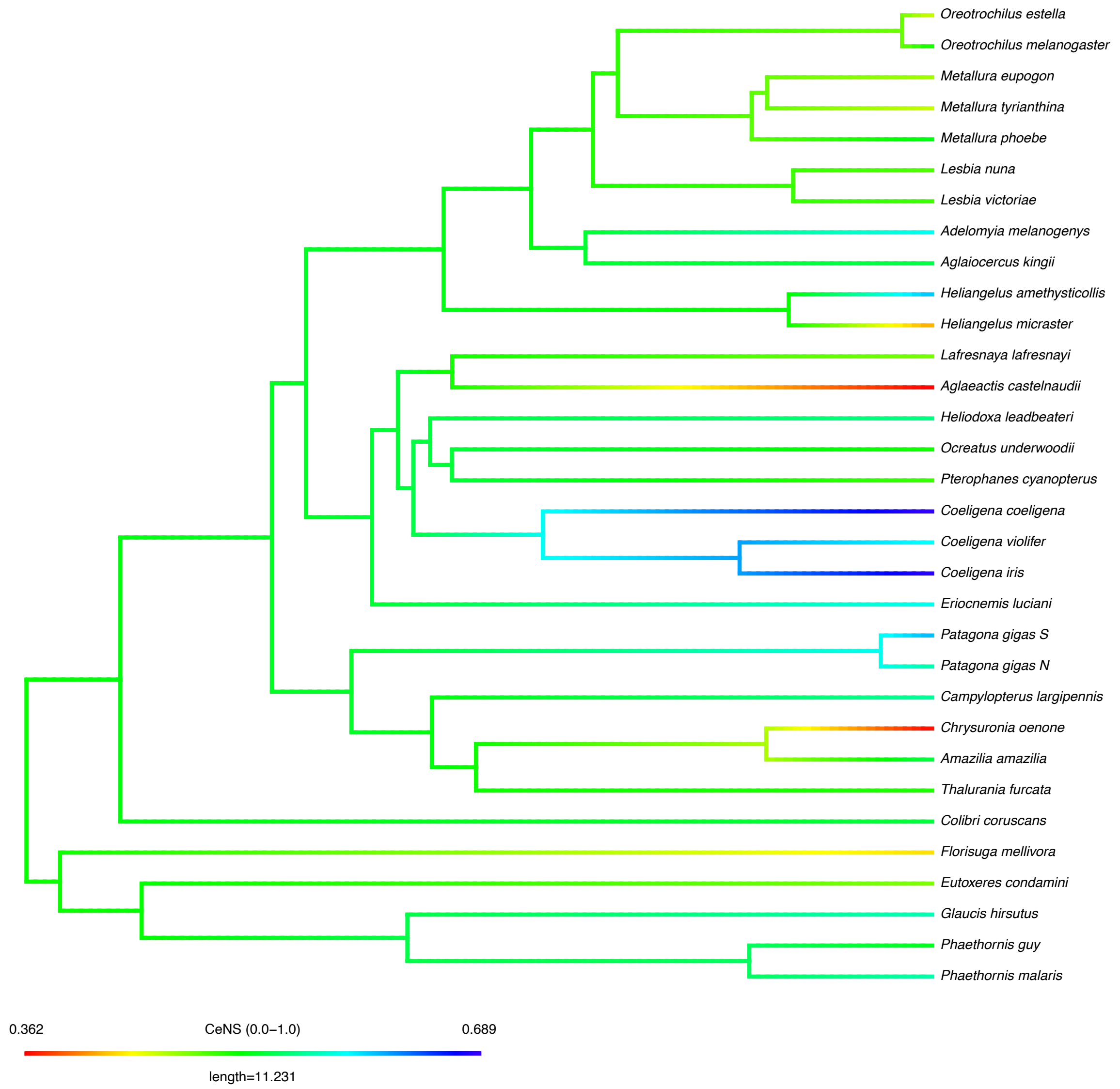


**Figure S4.** Continuous trait map of *CeNS* values showing modest phylogenenetic signal ($\lambda$ = 0.44, $p$ = 0.24; K < 0, $p$ = 1.0). *CeNS* ranged from 0 to 1, with higher values indicating proportionally greater contribution of TRBC than MCV in increasing [Hb]; similarly, lower values indicate proportionally greater contributions of MCV than TRBC in increasing [Hb]. Each *CeNS* value was mapped as a continuous trait using the contMap() function in *phytools* [1]. The tree is modified from McGuire et al. [2] and is pruned to the list of species in the *CeNS* modeling dataset (see Methods). Each species represented has ≥ 8 individuals sampled.

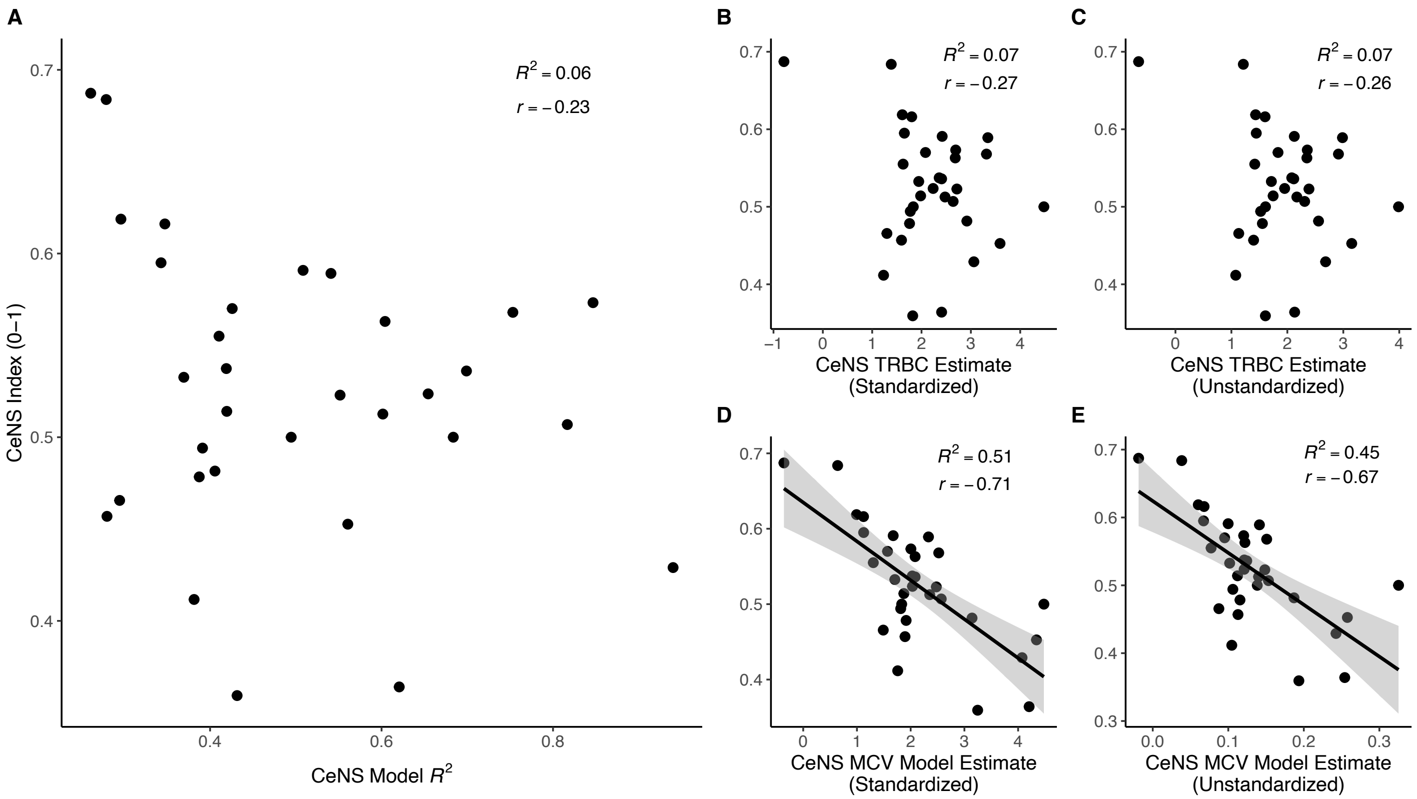


**Figure S5.** *CeNS* is not predicted by model fit **(A)** or TRBC estimates **(B–C),** but it is significantly predicted by MCV estimates (**D–E**; $p < 0.001$, linear model). TRBC and MCV model estimate results are consistent across standardized **(B, D)** and unstandardized **(C, E)**

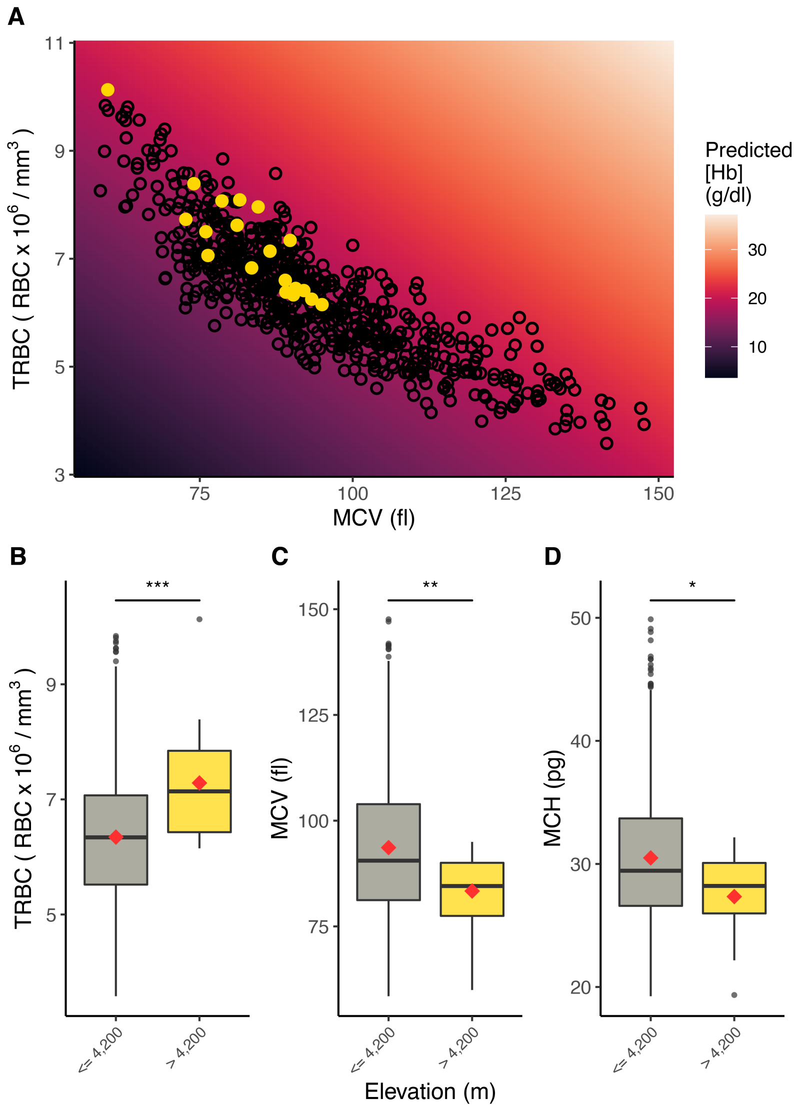


**Figure S6.** Blood values differ significantly at 'extreme' high elevations. **(A)** Total red blood cell count (TRBC) is significantly higher ≥4,200 m ($p < 0.001$). TRBC increases by 530,000 cells per µl with each 100-meter increase in elevation ≥4,200 m. **(B)** Mean cell volume (MCV) is significantly lower ≥4,200 m ($p$=0.010). MCV decreases by 5.52 fl with each 100-meter increase in elevation ≥4,200 m. **(C)** Mean cell hemoglobin content (MCH) is significantly lower ≥4,200 m ($p$=0.020). MCH decreases by 2.37 pg with each 100-meter increase in elevation ≥4,200 m. Data were analysed by subsetting the individual-level dataset by sampling elevation ("<4,200 m" or "≥4,200 m"), consistent with notable trends in the data (Figure 2). Significance and direction of relationships were assessed with one-way ANOVAs. Rates of increase and decrease were calculated using regression coefficients generalized linear models.

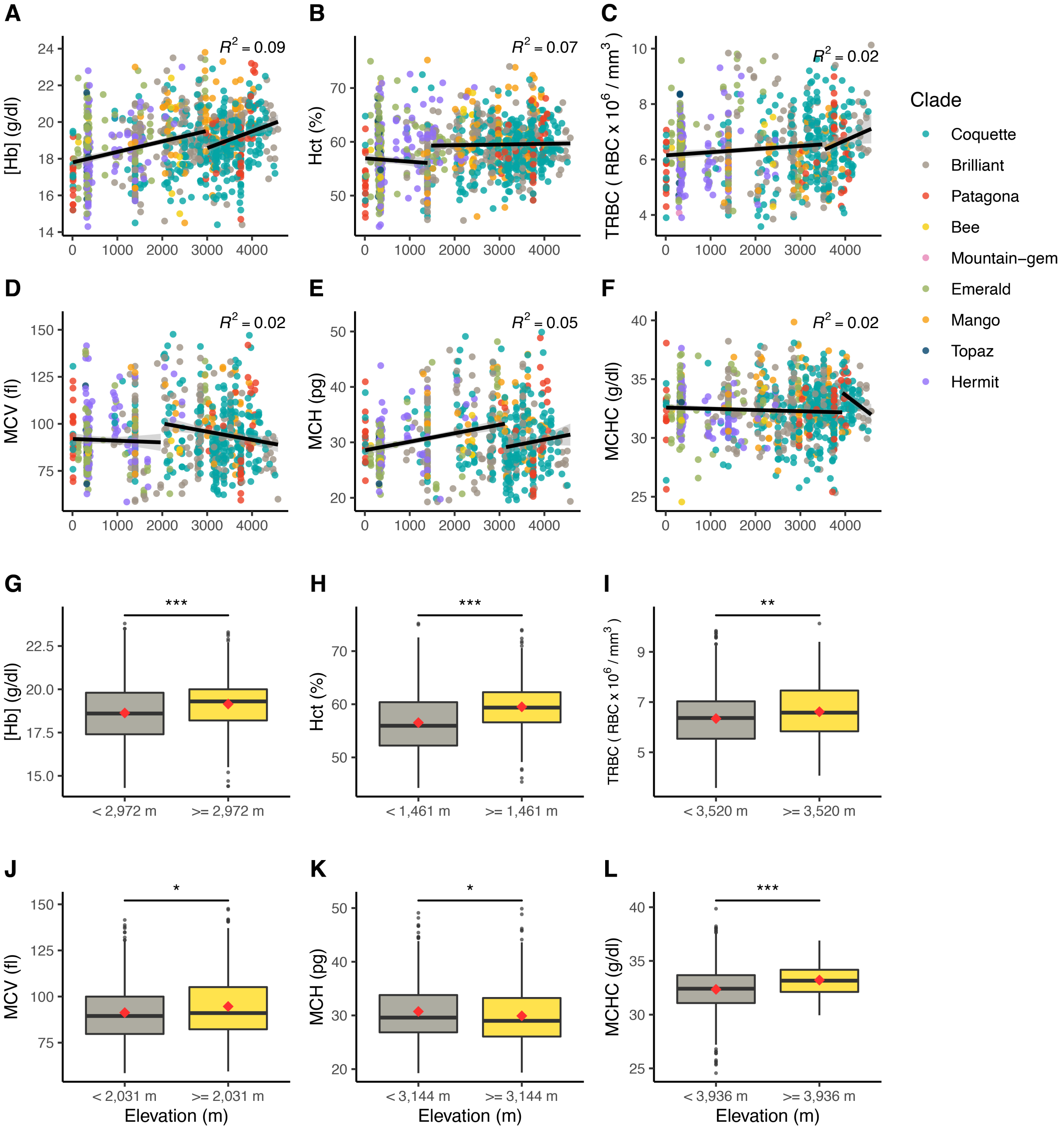


**Figure S7.** Relationship between elevation and the six studied blood parameters (hemoglobin concentration ([Hb]), hematocrit (Hct), total red blood cell count (TRBC), mean cell volume (MCV), mean cell hemoglobin content (MCH), and mean cell hemoglobin concentration (MCHC). Colors correspond to clades in Figure 1. **(A–F)** Within species comparisons, wherein each point represents a value from a single individual. Trend lines, 95% confidence intervals, and $R^2$ values are reported from piecewise models of elevation regressed on each predictor; in all cases, piecewise models fit better than simple linear regression models. **(A)** [Hb] increases by

0.06 g/dl and 0.09 g/dl for every 100-m increase in elevation below and above the breakpoint at 2,972 m, respectively. **(B)** Hct decreases by 0.06 and 0.01 percentage points for every 100-m increase in elevation below and above the breakpoint at 1,461 m, respectively. **(C)** TRBC increases by 10,000 cells per µl and by 70,000 cells per µl with each 100-meter increase below and above the breakpoint at 3,520 m, respectively. **(D)** MCV decreases by 0.09 fl and 0.44 fl for every 100-m increase in elevation below and above the breakpoint at 2,031, respectively. **(E)** MCH increases by 0.15 pg and 0.16 pg for every 100-m increase in elevation below and above the breakpoint at 3,144 m, respectively. **(F)** MCHC decreases by 0.01 g/dl and 0.29 g/dl with every 100-m increase in elevation below and above the breakpoint at 3,936 m, respectively. Rates of increase and decrease for each parameter were calculated using regression coefficients generalized linear models. **(G–L)** All blood traits differ significantly below and above breakpoint thresholds estimated from piecewise regression models (all $p < 0.001$). Model-estimated breakpoint thresholds for each blood trait are presented on the x-axis of each box plot. Significance and direction of relationships were assessed with one-way ANOVAs. Data were analysed by subsetting the individual-level dataset by model-estimated breakpoints (“<breakpoint” or “>=breakpoint”).

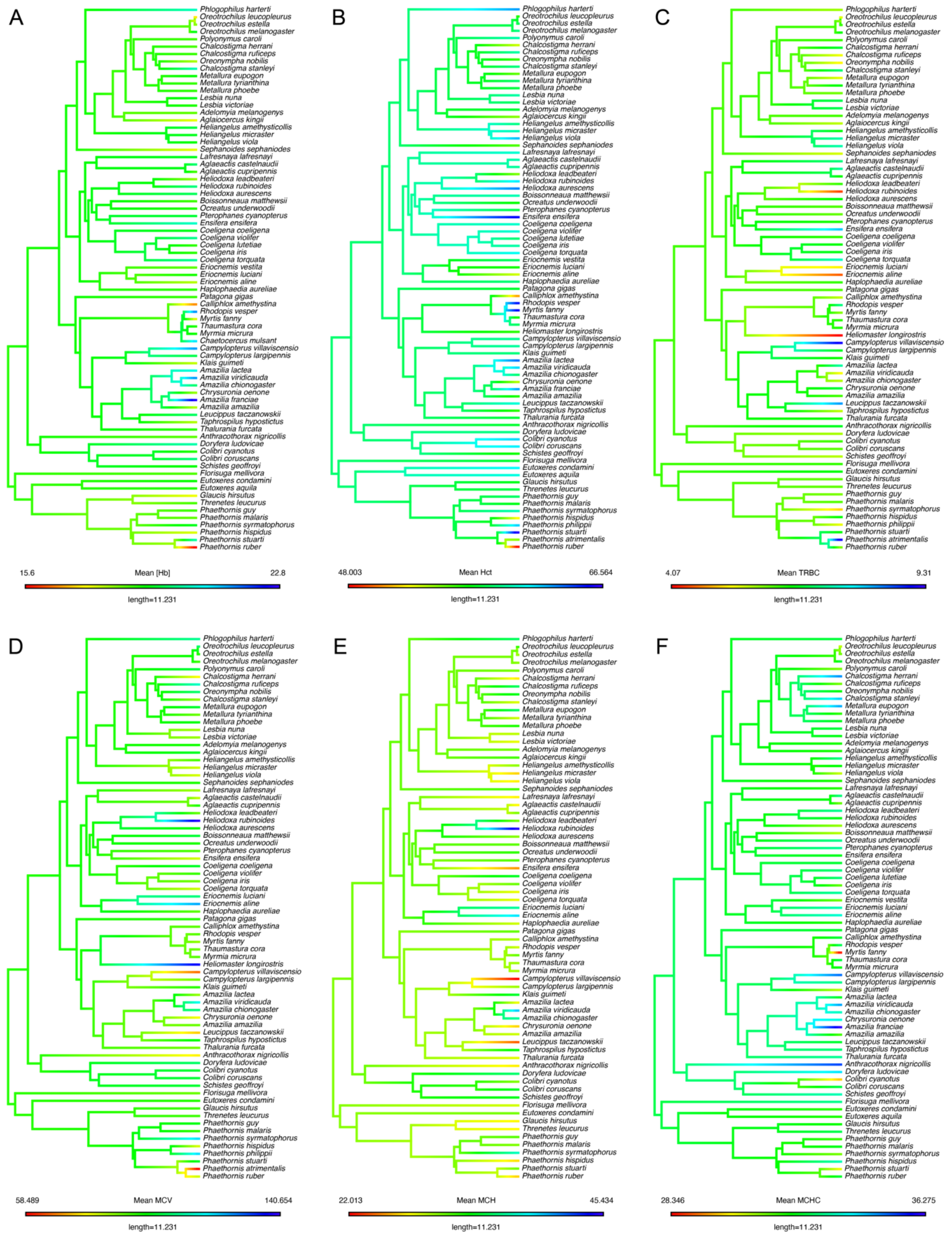


**Figure S8.** Continuous trait maps of mean trait values for each blood parameter based on within species data. **(A)** hemoglobin concentration ([Hb]); **(B)** hematocrit (Hct); **(C)** total red blood cell count (TRBC); **(D)** mean cell volume (MCV); **(E)** mean cell hemoglobin (MCH); **(F)** mean cell hemoglobin concentration (MCHC). The mean trait value for each hummingbird bird species was mapped as a continuous trait using the contMap() function in *phytools* [1]. The tree is modified from McGuire et al. [2] (see Methods) and pruned to the list of species in each blood data subset. In several cases, high signal (i.e., bright red or bright blue tips) is related to low species sample size ($n < 3$; Figure 1); removing these species and re-running tests of phylogenetic signal did not affect model results. Median trait values yield consistent results to mean trait values.

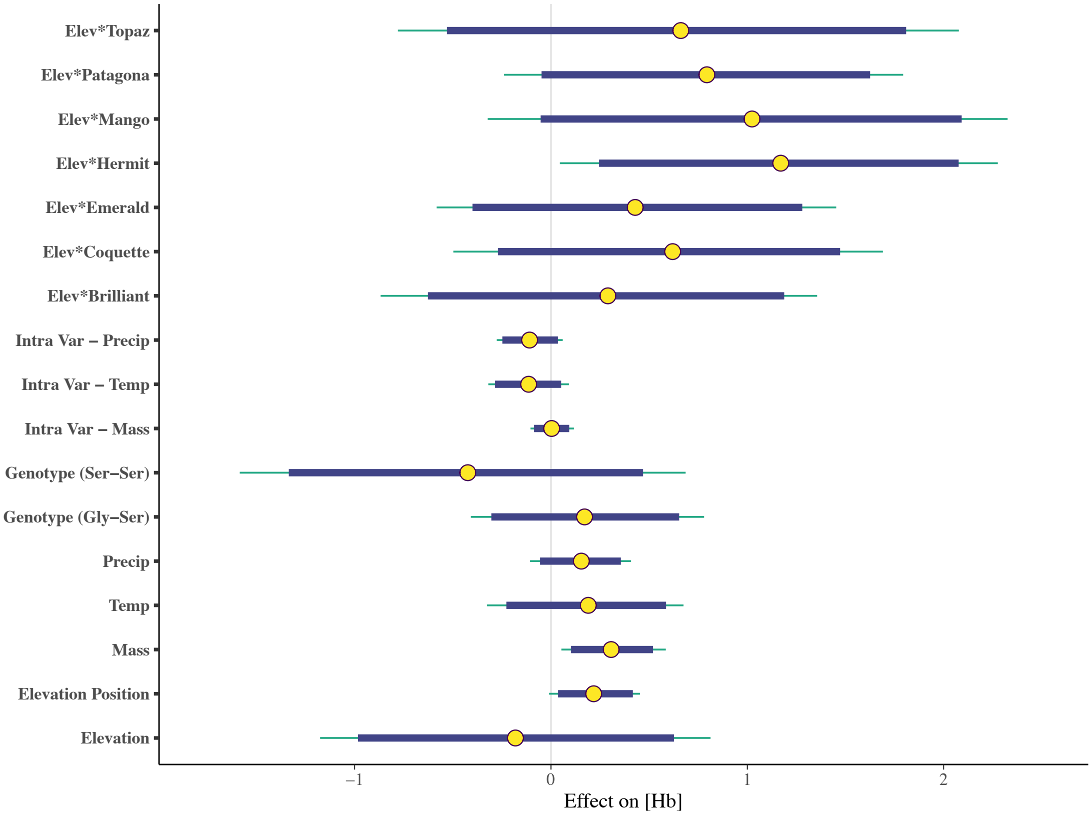


**Figure S9.** We found no clade*elevation interactions in our data with the exception of the interaction of elevation and the Hermit clade in the [Hb] model set. Circles indicate parameter estimates ($\beta$s), and thick and thin bars illustrate 89% and 95% credible intervals, respectively. The positive effect of elevation*Hermit on [Hb] and lack of 95% credible intervals overlap with zero suggests that Hermits have a notably steeper [Hb] slope than other clades.

**Table S1.** Important terms related to blood-$O_2$ carrying capacity, with relevant abbreviations and definitions. Equations for secondary blood indices are from Campbell & Ellis [3].

| Term | Abbreviation | Definition |
|---|---|---|
| Hemoglobin | Hb | Blood protein that transports oxygen to the cells |
| Hemoglobin concentration | [Hb] | Amount of hemoglobin relative to total volume in cells |
| Hematocrit | Hct | Proportion of the total volume of blood that is composed of red blood cells; synonymous with packed cell volume (PCV) and typically expressed as a percentage. |
| Mean cellular hemoglobin | MCH | Mean amount of hemoglobin in each red blood cell (pg); also called mean corpuscular hemoglobin in the literature. Expressed as *([Hb] / TRBC) x 10*. |
| Mean cellular hemoglobin concentration | MCHC | Mean concentration of hemoglobin per cell (gm/dl). Expressed as *([Hb] / Hct %) x 100* |
| Mean cell volume | MCV | Mean size of erythrocytes (fl); also called mean corpuscular volume or red cell volume in the literature. Expressed as *(Hct % / TRBC) x 10*. |
| Packed cell volume | PCV | Percentage of whole blood composed of erythrocytes (red blood cells); synonymous to hematocrit (Hct) and typically expressed as a percentage |
| Total red blood cell count | TRBC | Total number of red blood cells in millions per cubic millimeter (cumm or $mm^3$) of blood, or RBC x $10^6/mm^3$. One cubic millimeter is equivalent to one microliter (µl). Number of cells per microliter can be calculated by multiplying TRBC x $10^6$. |

**Table S2 .** Summary statistics for the individual-level dataset (*n*=1,217). For each parameter, we provide sample size of the data subset (*n*), mean, standard deviation, range (minimum and maximum values), and coefficient of variation.

| Parameter (units) | *n* | Mean | Standard Deviation | Minimum | Maximum | Coefficient of Variation |
|---|---|---|---|---|---|---|
| [Hb] (g/dl) | 988 | 18.84 | 1.65 | 14.3 | 23.8 | 0.09 |
| Hct (%) | 1086 | 58.54 | 5.15 | 44.25 | 75.2 | 0.09 |
| TRBC (RBC x $10^6/mm^3$) | 748 | 6.37 | 1.16 | 3.58 | 10.13 | 0.18 |
| MCV (fl) | 732 | 93.36 | 17.1 | 58.49 | 147.62 | 0.18 |
| MCH (pg) | 703 | 30.4 | 5.64 | 19.25 | 49.88 | 0.19 |
| MCHC (g/dl) | 924 | 32.44 | 2.12 | 24.56 | 39.86 | 0.07 |

**Table S3.** Comparison of Bayesian models to evaluate within- and among-species variation in the 6 studied blood traits: hemoglobin concentration ([Hb]), hematocrit (Hct), total red blood cell count (TRBC), mean cell volume (MCV), mean cell hemoglobin (MCH), and mean cell hemoglobin concentration (MCHC). Only within species models included the species grouping variable and only among species TRBC, MCV, MCH, and MCHC models included a quadratic component (elev$^2$). Reduced models included the set of predictors from full models whose 95% credible intervals did not overlap zero. Dashes indicate that a reduced model was not present (i.e., all credible intervals from full models overlapped zero). Model fit was assessed using the widely applicable information criterion (WAIC) and leave-one-out cross-validation (LOO), which were consistent. The difference between the best fit model and each other model in the set is shown as ΔELPD with difference in standard error (ΔSE) in parentheses (LOOIC values are calculated as –2*ELPD, which converts ELPD to a deviance scale). ΔELPD < 4 indicates that differences between two models are small.

| | **Model description** | | | | | | **Response** | | | | | |
|---|---|---|---|---|---|---|---|---|---|---|---|---|
| | **Predictors** | | | | | **Grouping Variable** | **[Hb] ΔELPD (ΔSE)** | **Hct ΔELPD (ΔSE)** | **TRBC ΔELPD (ΔSE)** | **MCV ΔELPD (ΔSE)** | **MCH ΔELPD (ΔSE)** | **MCHC ΔELPD (ΔSE)** |
| | **None (null)** | **All (full)** | **Reduced** | **All + elev$^2$** | **Reduced + elev$^2$** | **Species** | | | | | | |
| Within species | X | | | NA | NA | X | -29 (8.4) | -22.6 (8.2) | -3.9 (4.7) | -0.5 (1.0) | **0.0 (1.5)** | -11.5 (5.0) |
| | | X | | NA | NA | X | **0.0** | **0.0** | **0.0** | -0.9 (3.6) | -1.2 (3.1) | -0.1 (2.6) |
| | | | X | NA | NA | X | -3.3 (4.2) | -0.3 (2.8) | -0.1 (3.8) | **0.0** | **0.0** | **0.0** |
| | X | | | NA | NA | | -85.5 (13.1) | -70.5 (13.6) | -34.6 (9.3) | -24.9 (6.4) | -19.6 (6.5) | -39.4 (8.7) |
| | | X | | NA | NA | | -29.9 (7.6) | -29.9 (7.7) | -29.8 (7.7) | -26.5 (7.3) | -22.8 (6.7) | -29.3 (7.1) |
| | | | X | NA | NA | | -29.7 (8.8) | -26.9 (7.9) | -25.0 (8.2) | -23.3 (7.1) | -20.1 (6.5) | -29.9 7.1 |
| Among species | X | | | | | NA | -4.5 (3.9) | -2.3 (4.1) | **0.0** | **0.0** | -1.3 (2.2) | **0.0** |
| | | X | | | | NA | -0.1 (3.9) | **0.0** | -2.7 (2.6) | -3.6 (2.5) | -4.5 (3.5) | -5.4 (1.8) |
| | | | X | | | NA | **0.0** | 0.1 (2.0) | -0.5 (0.8) | – | – | – |
| | | | | X | | NA | NA | NA | -2.0 (2.5) | -2.4 (2.7) | -2.8 (2.1) | -4.5 (2.1) |
| | | | | | X | NA | NA | NA | – | – | **0.0** | – |

**Table S4.** Parameter estimates (*β*) and lower (LCI) and upper (UCI) 95% credible intervals from top Bayesian models for within and among species blood trait models. Bold indicates parameters of high importance for which 95% credible intervals did not overlap zero (Figure 4). Dashes indicate that a parameter was not present in a final model set. 'NA' indicates that a parameter was not included. For all models, Genotype: Gly-Gly was the reference category. A comparison of models across sets is presented in Table 1.

| | | **[Hb]** | | | **Hct** | | | **TRBC** | | | **MCV** | | | **MCH** | | | **MCHC** | | |
|---|---|---|---|---|---|---|---|---|---|---|---|---|---|---|---|---|---|---|---|
| | **Parameter** | **β** | **LCI** | **UCI** | **β** | **LCI** | **UCI** | **β** | **LCI** | **UCI** | **β** | **LCI** | **UCI** | **β** | **LCI** | **UCI** | **β** | **LCI** | **UCI** |
| **Within species** | Intercept | 18.84 | 18.45 | 19.23 | 58.65 | 57.47 | 59.87 | 6.51 | 6.14 | 6.86 | 94.29 | 92.20 | 96.58 | 30.70 | 30.03 | 31.42 | 32.51 | 32.25 | 32.76 |
| | Elevation | **0.52** | **0.16** | **0.9** | **1.87** | **0.71** | **3.01** | **0.41** | **0.08** | **0.74** | – | – | – | -0.63 | -1.60 | 0.40 | – | – | – |
| | Elevation position | 0.18 | -0.04 | 0.39 | **0.67** | **0.03** | **1.32** | 0.11 | -0.09 | 0.3 | – | – | – | – | – | – | – | – | – |
| | Mass | 0.21 | -0.02 | 0.42 | 0.48 | -0.2 | 1.14 | 0.04 | -0.15 | 0.24 | – | – | – | – | – | – | – | – | – |
| | Temperature | 0.28 | -0.12 | 0.7 | 1.19 | -0.04 | 2.51 | 0.37 | -0.01 | 0.75 | -1.08 | 0.85 | -2.79 | -0.81 | -1.79 | 0.24 | | | |
| | Precipitation | 0.07 | -0.17 | 0.31 | 0.2 | -0.51 | 0.95 | -0.08 | -0.29 | 0.12 | – | – | – | – | – | – | – | – | – |
| | Genotype: Gly-Ser | 0.16 | -0.37 | 0.69 | 0.01 | -1.62 | 1.64 | -0.32 | -0.86 | 0.21 | – | – | – | – | – | – | – | – | – |
| | Genotype: Ser-Ser | -0.23 | -1.22 | 0.73 | -0.7 | -4.02 | 2.31 | -0.07 | -1.05 | 0.89 | – | – | – | – | – | – | – | – | – |
| | Intra Var: Mass | 0.03 | -0.08 | 0.13 | **-0.44** | **-0.74** | **-0.14** | -0.06 | -0.16 | 0.04 | – | – | – | – | – | – | **0.20** | **0.07** | **0.34** |
| | Intra Var: Temp | -0.16 | -0.34 | 0.01 | -0.22 | -0.76 | 0.3 | -0.01 | -0.17 | 0.15 | – | – | – | – | – | – | **-0.23** | **-0.35** | **-0.10** |
| | Intra Var: Precip | -0.07 | -0.23 | 0.1 | -0.05 | -0.55 | 0.44 | 0.04 | -0.09 | 0.17 | – | – | – | – | – | – | – | – | – |
| | *Species grouping variable* | 0.13 | 0.06 | 0.23 | 0.13 | 0.06 | 0.22 | 0.21 | 0.11 | 0.33 | 0.13 | 0.06 | 0.24 | 0.12 | 0.05 | 0.22 | 0.15 | 0.07 | 0.25 |
| **Among species** | Intercept | 18.75 | 18.54 | 18.96 | 58.83 | 57.71 | 59.94 | 6.26 | 6.07 | 6.46 | 95.05 | 92.27 | 97.95 | 31.87 | 30.48 | 33.25 | 32.51 | 32.24 | 32.78 |
| | Elevation | **0.36** | **0.16** | **0.58** | **1.83** | **0.53** | **3.08** | – | – | – | – | – | – | 0.51 | -0.4 | 1.37 | – | – | – |
| | Elevation Quadratic | NA | NA | NA | NA | NA | NA | – | – | – | – | – | – | **-1.2** | **-2.28** | **-0.09** | – | – | – |
| | Mass | – | – | – | 0.64 | -0.12 | 1.41 | – | – | – | – | – | – | – | – | – | – | – | – |
| | Temperature | – | – | – | 0.9 | -0.71 | 2.54 | – | – | – | – | – | – | – | – | – | – | – | – |
| | Precipitation | – | – | – | -0.02 | -1.3 | 1.3 | – | – | – | – | – | – | – | – | – | – | – | – |
| | Genotype: Gly-ser | – | – | – | -0.83 | -2.67 | 1.02 | – | – | – | – | – | – | – | – | – | – | – | – |
| | Genotype: Ser-Ser | – | – | – | -2.12 | -5.37 | 1.24 | – | – | – | – | – | – | – | – | – | – | – | – |

**Table S5.** Correlations between pairs of blood parameters for the individual-level (unshaded, top half) and species-level (gray shaded, bottom half) datasets, respectively. Correlations for the individual-level dataset were calculated from 688 values due to removal of 'NA' values from the combined dataset. Likewise, correlations for species-level dataset were calculated from 667 observations due to exclusion of 'NA' observations and did not exclude distributional or sampling outliers.

| **Parameter** | [Hb] | Hct | TRBC | MCV | MCH | MCHC |
|---|---|---|---|---|---|---|
| [Hb] | – | 0.76 | 0.24 | 0.14 | 0.25 | 0.34 |
| Hct | 0.76 | – | 0.30 | 0.20 | 0.08 | -0.35 |
| TRBC | 0.24 | 0.30 | – | -0.85 | -0.85 | -0.08 |
| MCV | 0.14 | 0.20 | -0.85 | – | 0.94 | -0.09 |
| MCH | 0.25 | 0.08 | -0.85 | 0.94 | – | 0.25 |
| MCHC | 0.34 | -0.35 | -0.08 | -0.09 | 0.25 | – |
| [Hb] | – | 0.83 | 0.47 | -0.04 | 0.05 | 0.24 |
| Hct | 0.83 | – | 0.34 | 0.16 | 0.08 | -0.34 |
| TRBC | 0.47 | 0.34 | – | -0.85 | -0.84 | 0.19 |
| MCV | -0.04 | 0.16 | -0.85 | – | 0.96 | -0.33 |
| MCH | 0.05 | 0.08 | -0.84 | 0.96 | – | -0.05 |
| MCHC | 0.24 | -0.34 | 0.19 | -0.33 | -0.05 | – |

**Table S6.** Comparison of Bayesian models to assess within and among species variation in the importance of cell size versus number in increasing [Hb], and parameter estimates (*β*) and lower (LCI) and upper (UCI) 95% credible intervals from top (full) models. Each set included two models: an intercept-only null model, and a model with total red blood cell count (TRBC) and mean cell volume (MCV) as predictors. Within species models included a species grouping variable (i.e., "+ (1|species)"). Model fit was assessed using the widely applicable information criterion (WAIC) and leave-one-out cross-validation (LOO), respectively, which were consistent. The difference between the best fit model and each other model in the set is shown as ΔELPD with difference in standard error (ΔSE) in parentheses (LOOIC values are calculated as –2*ELPD, which converts ELPD to a deviance scale). Bold indicates parameters of high importance for which 95% credible intervals did not overlap zero (Figure 5). NA indicates that a parameter was not included.

| | **Model description** | | | **Response** | **βs (95% LCI–UCI)** | | |
|---|---|---|---|---|---|---|---|
| | **Predictors** | | **Grouping Variable (% variance explained)** | **[Hb] ΔELPD (ΔSE)** | **Intercept** | **TRBC** | **MCV** |
| | **None** | **All** | **Species** | | | | |
| Within species | X | | NA | -243.5 (19.1) | 18.84 (18.71-18.96) | NA | NA |
| | | X | 14% | 0.0 | 18.87 (18.70-19.03) | **2.05 (1.87-2.22)** | **1.92 (1.75-2.10)** |
| Among species | X | | NA | -21.5 (5.3) | 18.95 (18.67-19.21) | NA | NA |
| | | X | NA | 0.0 | 19.94 (18.77-19.12) | **1.45 (1.12-1.79)** | **1.41 (1.08-1.75)** |

**Table S7.** Comparison of Bayesian models to understand factors that contribute to variation in *CeNS*, a measure of the proportional contribution of TRBC to MCV in increasing [Hb] and parameter estimates (*β*) and lower (LCI) and upper (UCI) 95% credible intervals from the top *CeNS* model. Reduced models included the set of predictors from full models whose 95% credible intervals did not overlap zero (e.g., Figure 5A). "Elevation$^2$" indicates the quadratic elevation term. For all models, Genotype: Gly-Gly was the reference category.
Model fit was assessed using the widely applicable information criterion (WAIC) and leave-one-out cross-validation (LOO), which were consistent. The difference between the best fit model and each other model in the set is shown as ΔELPD with difference in standard error (ΔSE) in parentheses (LOOIC values are calculated as –2*ELPD, which converts ELPD to a deviance scale). ΔELPD < 4 indicates that differences between two models are small. Bolded *β* estimates indicate parameters of high importance from the top *CeNS* prediction model whose 95% credible intervals did not overlap zero (Figure 5A).

| **Model** | **ΔELPD** | **ΔSE** | **Top *CeNS* Prediction Model (elevation + elevation$^2$ + mass + wing loading)** | | | |
|---|---|---|---|---|---|---|
| | | | **Parameter** | **β** | **LCI** | **UCI** |
| elevation + elevation$^2$ + mass + wing loading | 0.0 | 0.0 | Intercept | 0.58 | 0.53 | 0.63 |
| elevation + elevation$^2$ + mass | -0.1 | 1.4 | Elevation | **-0.04** | **-0.07** | **-0.01** |
| elevation + elevation$^2$ + mass + wing loading + genotype | -1.5 | 0.9 | Elevation$^2$ | **-0.05** | **-0.09** | **-0.02** |
| null (intercept-only) | -3.1 | 3.5 | Mass | **0.04** | **0.01** | **0.06** |
| elevation + mass + wing loading | -3.1 | 2.9 | Wing Loading | -0.02 | -0.05 | 0.01 |
| elevation + elevation$^2$ | -3.4 | 2.8 | | | | |

**Table S8.** Results of phylogenetic signal tests (Blomberg's K and Pagel's λ) for intraspecific models (individual-level data), showing low phylogenetic signal across all blood traits of interest. For Blomberg's K, values close to or above one are thought to indicate strong phylogenetic signal. Values for Pagel's λ are bounded between 0 and 1, with values close to 0 indicating no phylogenetic signal and values close to 1 indicating strong phylogenetic signal. The weak or negligible phylogenetic signal present in these data is confirmed by visual examination of continuous trait maps (Figure S1).

| Trait | Blomberg's K | Blomberg's K *p*-value | Pagel's λ | LogL(λ) | LR(λ =0) | Pagel's λ *p*-value |
|---|---|---|---|---|---|---|
| **Hb** | 0.142 | 0.824 | 0.000 | -117.240 | 0.000 | 1 |
| **Hct** | 0.172 | 0.721 | 0.000 | -199.735 | -0.002 | 1 |
| **TRBC** | 0.153 | 0.783 | 0.000 | -96.119 | -0.002 | 1 |
| **MCV** | 0.208 | 0.640 | 0.000 | -291.408 | -0.002 | 1 |
| **MCH** | 0.264 | 0.395 | 0.000 | -191.235 | -0.002 | 1 |
| **MCHC** | 0.209 | 0.641 | 0.000 | -122.802 | -0.001 | 1 |

**REFERENCES**

1. Revell LJ. phytools: an R package for phylogenetic comparative biology (and other things). Methods Ecol Evol. 2012;3: 217–223. doi:10.1111/j.2041-210X.2011.00169.x
2. McGuire JA, Witt CC, Remsen J V, Corl A, Rabosky DL, Altshuler DL, et al. Molecular phylogenetics and the diversification of hummingbirds. Curr Biol. 2014;24: 1–7. doi:10.1016/j.cub.2014.03.016
3. Campbell TW, Ellis CK. Avian and Exotic Animal Hematology and Cytology. Blackwell Publishing Inc.; 2007. doi:10.1017/CBO9781107415324.004
